# Differences in clamp loader mechanism between bacteria and eukaryotes

**DOI:** 10.1101/2023.11.30.569468

**Authors:** Jacob T. Landeck, Joshua Pajak, Emily K. Norman, Emma L. Sedivy, Brian A. Kelch

## Abstract

Clamp loaders are pentameric ATPases that place circular sliding clamps onto DNA, where they function in DNA replication and genome integrity. The central activity of a clamp loader is the opening of the ring-shaped sliding clamp, and the subsequent binding to primer-template (p/t)-junctions. The general architecture of clamp loaders is conserved across all life, suggesting that their mechanism is retained. Recent structural studies of the eukaryotic clamp loader Replication Factor C (RFC) revealed that it functions using a crab-claw mechanism, where clamp opening is coupled to a massive conformational change in the loader. Here we investigate the clamp loading mechanism of the *E. coli* clamp loader at high resolution using cryo-electron microscopy (cryo-EM). We find that the *E. coli* clamp loader opens the clamp using a crab-claw motion at a single pivot point, whereas the eukaryotic RFC loader uses motions distributed across the complex. Furthermore, we find clamp opening occurs in multiple steps, starting with a partly open state with a spiral conformation, and proceeding to a wide open clamp in a surprising planar geometry. Finally, our structures in the presence of p/t-junctions illustrate how clamp closes around p/t-junctions and how the clamp loader initiates release from the loaded clamp. Our results reveal mechanistic distinctions in a macromolecular machine that is conserved across all domains of life.

## INTRODUCTION

Sliding clamps and clamp loaders are integral components of the DNA replication machinery throughout all known life. Sliding clamps are ring-shaped protein complexes that encircle DNA in order to tether proteins to the genome^1^. Because the sliding clamp is a closed ring in solution, it needs to be opened and placed onto DNA by a separate ATPase complex called the clamp loader. Sliding clamps and clamp loaders are necessary for numerous aspects of DNA replication and repair, thus these protein complexes are important for human health^2^.

All clamp loaders are pentameric ATPases of the AAA+ family (<u>A</u>TPases <u>a</u>ssociated with various cellular <u>a</u>ctivities)^3,4^. In the presence of ATP, clamp loaders bind to the sliding clamp and open the ring (**Fig. 1C**)^5–7^. This open binary complex can then attach to a primer-template (p/t)-junction, which triggers ATPase activity^8–11^. ATP hydrolysis results in closure of the sliding clamp and ejection of the clamp loader^12–15^. Therefore, clamp loaders can be thought of as protein-remodeling switches^16–18^, in contrast to most AAA+ proteins that act as processive motors^19^.

**Figure 1.**
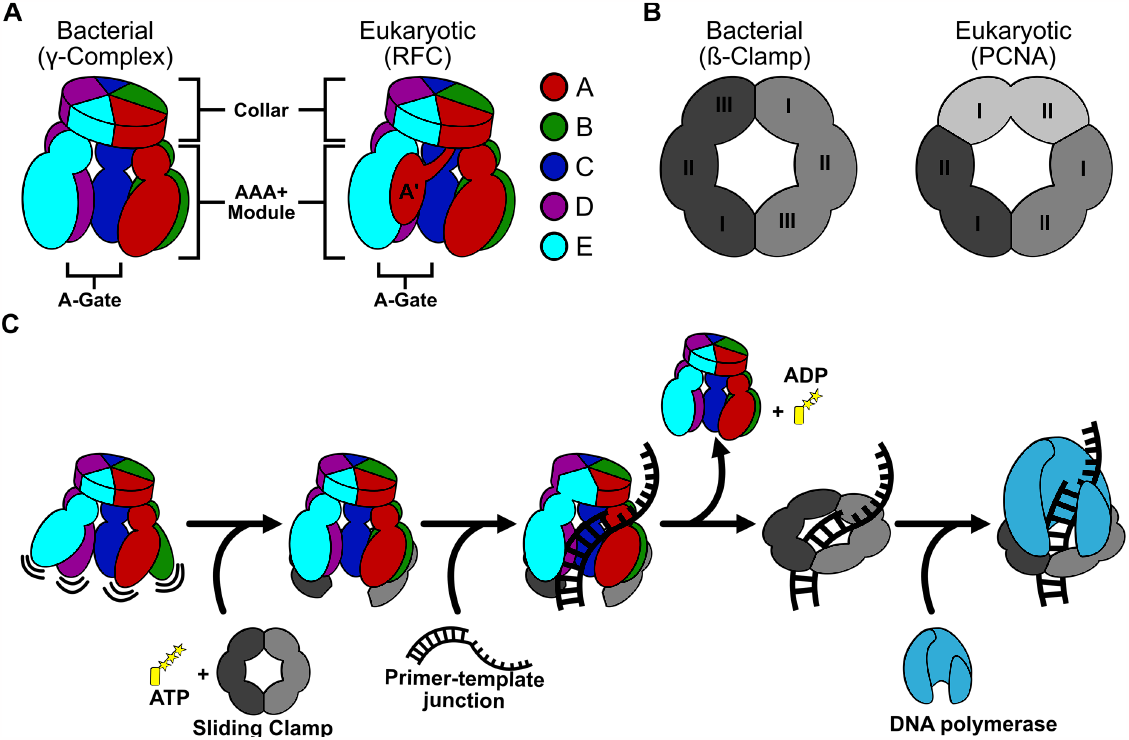
General architecture and mechanism of clamp loaders and sliding clamps. **A)** Comparison of bacterial and eukaryotic clamp loaders. Clamp loaders are pentamers with each subunit consisting of a AAA+ module and a collar domain. The “A-gate”, a gap between the A and E subunits, is where p/t-junctions enter. The A’ domain is unique to eukaryotic clamp loaders. **B)** Comparison of bacterial and eukaryotic sliding clamps. The bacterial sliding clamp is a homodimer where each subunit has three domains. The eukaryotic sliding clamp is a homotrimer, where each subunit has two domains. **C)** Summary of the clamp loading mechanism. Clamp loaders first bind ATP, then the sliding clamp. The clamp loader then opens the sliding clamp and the complex binds a p/t-junction. After ATP hydrolysis and clamp closure, the clamp loader releases the sliding clamp. The loaded sliding clamp can then be used by DNA replication and repair factors.

Despite the overall conservation of this machinery, there are key differences in subunit stoichiometry and structure across different domains of life. While all sliding clamp proteins have six domains arranged into a ring, bacterial sliding clamps (also known as ß-clamps) are dimers of three domains^1^, and eukaryotic clamps (Proliferating Cell Nuclear Antigen or PCNA) are trimers of two domains (**Fig. 1B**)^20,21^. Moreover, clamp loader subunit stoichiometry is distinct between eukaryotes and bacteria, despite the fact that they are all pentamers (with the subunits designated A through E). Eukaryotic loaders have five distinct proteins (RFC1 through RFC5), while bacteria have three proteins that are arranged in a 3:1:1 ratio (γ_3_δδ’, referred to as γ-complex hereafter; **Fig. 1A**). Moreover, the A subunit of eukaryotic clamp loaders contains a region called the A’ domain that is not present in bacterial clamp loaders^22–25^. It remains unclear how or if these differences in composition and structure affect the mechanism of clamp loading.

Structural studies of clamp loaders in action have revealed important insights into their mechanism. Numerous structures indicate that the eukaryotic loader (Replication Factor C or RFC) binds to its cognate clamp PCNA in a closed form, and then opens the clamp with a concomitant opening of the RFC structure^22,26–30^. The key motion is the opening of the “A-gate” at the A subunit, which allows p/t-DNA to enter into the complex. This conformational change has been described as a “crab-claw” mechanism for clamp opening. However, earlier fluorescence studies of the *E. coli* γ-complex predicted that the crab-claw mechanism does not apply^31^. In RFC, the A’ domain makes important interactions that control the opening of the A-gate^27^, but bacterial clamp loaders lack an A’ domain^32^. These observations raise the possibility that bacterial and eukaryotic clamp loaders open clamps using different mechanisms.

Here we describe a series of structures of the *E. coli* clamp loader in the process of placing the clamp onto a p/t-junction. These structures show that the bacterial clamp loader also uses a crab-claw mechanism, but it is quite different from what we observed in the eukaryotic system. The γ-complex binds to the ß-clamp and, through a multi-step process, opens the clamp into a planar configuration. The planar opening requires a single pivoting motion in between the B and C subunits, which disrupts the interfaces at the ATPase active site, preventing ATP hydrolysis. Subsequent binding of a p/t-junction activates the ATPase through tight interactions between adjacent AAA+ modules. Our results reveal a distinction in clamp loading between bacteria and eukaryotes, potentially opening the door for specific inhibition of bacterial clamp loaders as a new target for antibiotic development.

## RESULTS

### Structures of *E. coli* clamp loader opening the ß-clamp

We reconstituted the clamp opening reaction using a purified recombinant sliding clamp (ß) and clamp loader (γ-complex) (**Fig. S1A**). The reconstituted clamp loader is active because it displays the expected response to ATP and p/t-DNA (**Fig. S1B**). As seen in previous studies^5,33^, ß-clamp binding suppresses ATPase activity, but the presence of clamp and p/t-DNA synergistically activates ATPase activity.

To reveal how the *E. coli* clamp loader opens its clamp, we used single particle cryo-electron microscopy (cryo-EM) to determine structures of various clamp opening intermediates (**Fig. S1C**). In order to suppress the background ATPase activity in our cryo-EM samples, we replaced ATP with either the slowly hydrolysable ATP analog ATPγS or the non-hydrolysable analog ADP•BeF_x_. From these two cryo-EM datasets, we obtained three distinct reconstructions of the sliding clamp/clamp loader complex.

#### Clamp loader bound to closed clamp

In the presence of ATPγS, we obtained an ∼8 Å resolution reconstruction of the clamp loader bound to a closed clamp (**Fig. 2A; Fig. S2**). Despite the modest resolution of the reconstruction, we can unambiguously identify the location and orientation of all the protein domains of both the clamp loader and sliding clamp. We modeled these domains as rigid bodies, followed by Molecular Dynamics Flexible Fitting^34,35^ to adjust the domains while preventing steric clashes (**Fig. S3A**).

**Figure 2.**
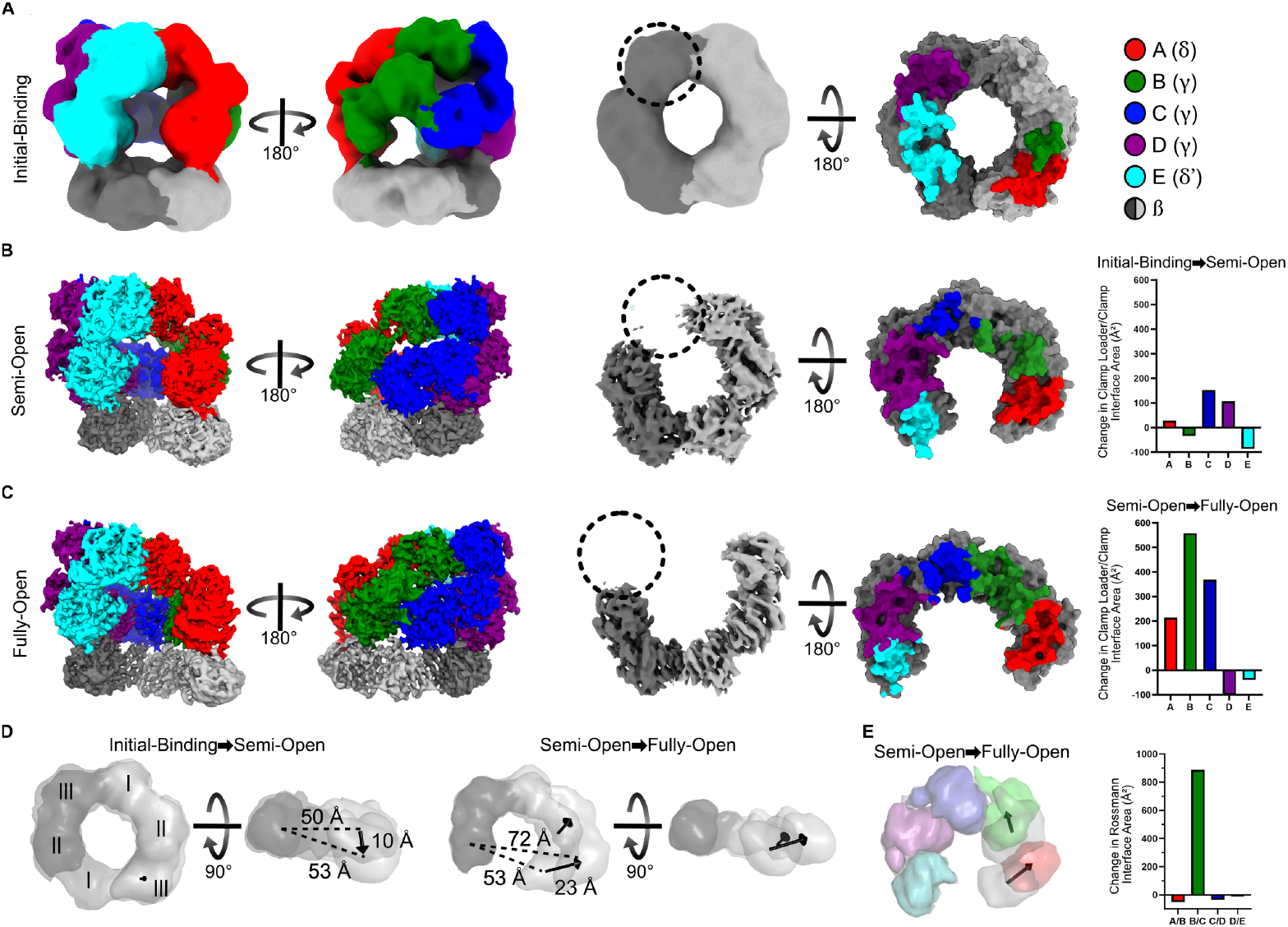
Structures of the clamp loader/sliding clamp complex throughout the process of clamp opening. **A-C)** The clamp loader/sliding clamp complex in the Initial-binding (A), Semi-Open (B), and Fully-Open (C) states. Left, cryo-EM maps in two views. Center, bottom view of the cryo-EM maps of bound sliding clamps. Right, a top view of the sliding clamp with residues that contact the clamp loader colored according to the contacting clamp loader subunit. The change in clamp/clamp loader interface area between states is quantified on the far right. There is no recognizable density for domain I of subunit II (indicated with a dashed circle) of the sliding clamp in the Semi-Open and Fully-Open states. **D)** Conformational changes of the sliding clamp upon opening. Models of the Initial-Binding, Semi-Open, and the Fully-Open states are displayed as low contour surfaces. Sliding clamps were aligned on subunit II and displacements of domains within subunit I are shown as vectors. We quantify clamp opening using the distance between the centers of mass of domain II of subunit II and domain III of subunit I (dashed lines). **E)** Conformational changes of the clamp loader between the Semi-Open and Fully-Open states. The Rossmann domains of the Semi-Open state (gray) and the Fully-Open state (color) are displayed as low contour surfaces. The structures are aligned on the Rossmann domain of subunit E, and the displacements of the centers of mass of the Rossmann domains of subunits A and B are shown as vectors. Subunits C, D, and E move less than 3 Å. The change in interface area between adjacent Rossmann domains is quantified to the right.

We observe that the γ-complex contacts the sliding clamp primarily through the A, D, and E subunits; the B and C subunits make minimal contact with the surface of the clamp (**Fig. 2A**). Furthermore, the AAA+ modules of the B and C subunits make minimal contact with each other (**Fig. S3B**). As inter-subunit contacts are necessary to hydrolyze ATP, this separation between subunits likely inhibits ATPase activity. Because the clamp is closed in this state, we attribute this conformation as the Initial-Binding complex.

#### Clamp Loader bound to Semi-Open or Fully-Open Clamp

In micrographs of the sample containing ADP•BeF_x_, we observe two different conformations of the clamp loader/sliding clamp complex, both with the clamp open (**Fig. S4**). However, the degree of opening is drastically different between these two states. In the first state, the clamp is barely opened and adopts a very tight spiral conformation (overall resolution = 3.9 Å; **Fig. 2B**). In the second state, the sliding clamp is opened into a wide, planar conformation (overall resolution = 3.0 Å; **Fig. 2C**). Therefore, we designate these two states as the Semi-Open and the Fully-Open states, respectively.

Unlike the Initial-Binding complex, in the Semi-Open state all five subunits of the clamp loader contact the sliding clamp (**Fig. 2B**). These additional contacts between the clamp loader and sliding clamp help stabilize the initial opening of the sliding clamp in a spiral conformation. Interface contacts between the A and B AAA+ modules and the C and D AAA+ modules are strong, while contacts between the B and C AAA+ modules remain minimal.

As in the Semi-Open state, all five subunits of the clamp loader also make contact with the sliding clamp in the Fully-Open state. The primary conformational change between these two states is a dramatic increase in interfacial contacts between the B and C subunits. This is accomplished by a single pivoting motion at a point between the B and C subunits (**Fig. 2E**). Because the B and C subunits are contacting the sliding clamp, this pivoting motion imparts an equivalent pivoting motion within subunit I of the sliding clamp, resulting in the wide planar opening (**Fig. 2D**). The first ß-clamp subunit undergoes a substantial change (C_α_ RMSD ∼3.0 Å), while the second ß-clamp subunit is essentially unchanged (C_α_ RMSD ∼0.7 Å). These motions pull the first subunit ‘upward’ into a flat “C” shape, rather than the spiral seen in structures of T4 phage and yeast clamp loading complexes^16,27–29^. A planar conformation of an open sliding clamp has been recently observed in the structure of the 9-1-1 clamp bound to the Rad24-RFC loader and resected DNA^23,24^.

In both the Semi- and Fully-Open states, there is no significant density for Domain I of the second subunit of the ß-clamp, which is at the site of ring opening (**Fig. 2B,C**). This is the only domain not directly held by a clamp loader subunit. All of the other domains of the ß-clamp are clearly observable, including Domain II directly adjacent to the missing domain. This indicates that the entirety of Domain I is too mobile to be clearly observed. This is consistent with hydrogen-deuterium exchange data, which indicates that this domain of the *E. coli* ß-clamp unfolds upon opening^36,37^. The lack of density for Domain I makes it challenging to measure the true width of the ß-clamp opening. Instead, we measure the distance between the centers of mass of Domain III of the first subunit and Domain II of the second (**Fig. 2D**).

### DNA binding activates ATP hydrolysis and closes the clamp

We next investigated how the *E. coli* clamp loader binds to p/t-junctions. Bacterial clamp loaders typically load onto RNA-primers at each Okazaki fragment *in vivo*^*38*^, but most *in vitro* measurements of structure or activity have been performed using DNA-primed p/t-junctions. Thus, we determined structures of the *E. coli* γ-complex and ß-clamp bound to both RNA-primed and DNA-primed p/t-junctions. To prevent hydrolysis of the bound nucleotide, we used the non-hydrolyzable ATP mimic ADP•BeFx for both samples. We determined the structure of the γ-complex bound to an open ß-clamp and an RNA-primed p/t-junction at 3.2 Å overall resolution (**Fig. 3C; Fig. S5**). The dataset using the DNA-primed p/t-junction resulted in four conformations of the ternary complex after 3D classification (**Fig. S6**). The first two are open states that differ in their collar domains (Open-DNAp/t and Altered-Collar), while the latter two are closed states that differ in their interactions between the loader and the clamp (Closed-DNA1 and Closed-DNA2).

**Figure 3.**
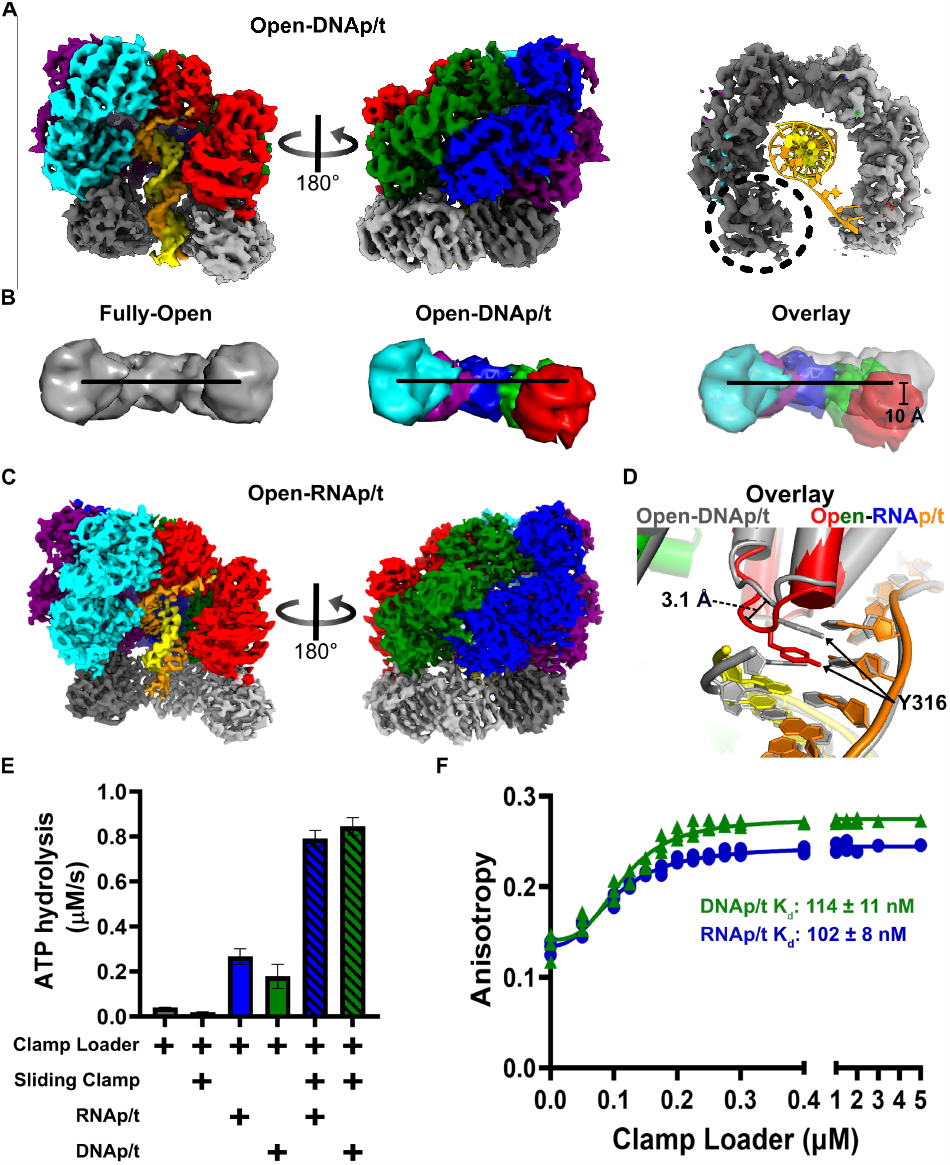
Clamp loader/sliding clamp complex bound to DNAp/t or RNAp/t. **A)** Cryo-EM map of the Open-DNAp/t complex. The clamp loader subunits are colored as before. The template strand is orange and the primer strand is yellow. The 3D reconstruction of the Open-DNAp/t complex has density for all six domains of the sliding clamp. **B)** The p/t-junction bound complex adopts a helical conformation. Models of the Rossmann domains of the Fully-Open (gray) and Open-DNAp/t (color) states are displayed as low contour surfaces. The black line displayed on all models connects the center of masses of the Rossmann domains of the A and E subunits of the Fully-Open state. The models were aligned on the E subunit. The center of mass of the A subunit shifts ∼10 Å after p/t-junction binding. **C)** Cryo-EM map of the Open-RNAp/t complex. **D)** The Open-DNAp/t (gray) and Open-RNAp/t (color) aligned on the collar of the A subunit. The DNA primer (gray) shifts registry, reaching higher into the central chamber compared to the RNA primer (yellow). This is accommodated by a 3.1 Å Cα shift in separation pin Y316, which stacks with the last base of the primer. **E)** Steady-State ATPase activity of the clamp loader stimulated by RNAp/t or DNAp/t. **F)** The affinity of the clamp loader for RNAp/t or DNAp/t measured by anisotropy of TAMRA-labeled template DNA.

#### Open-DNAp/t (overall resolution 3.7 Å)

We first discuss the next on-pathway state where p/t-DNA binds inside the clamp loader and clamp, causing both to constrict around DNA **(Fig. 3A)**. The primary motion is pivoting of the A and B subunits out-of-plane into a spiral. The AAA+ modules of the γ-complex and the sliding clamp adopt a helical arrangement that matches the helical symmetry of DNA **(Fig. 3B, Fig. S7B)**, similar to other structures of clamp loaders bound to p/t-DNA^16,27,39^. This constriction results in the A-gate closing by ∼5 Å, measured by the centers of mass of domains A and E. The clamp then closes slightly, by 11 Å **(Fig. S7A)**. Clamp closure is primarily driven by relaxation of ß-clamp subunit I into a conformation that is more like that of the closed form (C_α_-RMSD_FullyOpen_vs_Closed_ ∼3.3 Å versus C_α_-RMSD_DNA-Open_vs_Closed_ ∼1.8 Å).

#### Open-RNAp/t (overall resolution 3.2 Å)

We find that primer composition (RNA vs. DNA) has very little impact on the conformation of the complex. The C_α_ RMSD between the Open-RNAp/t structure and that of the Open-DNAp/t is only ∼0.7 Å **(Fig. S7D)**. Both structures show that the duplex region of the p/t-junction adopts A-form geometry as expected^39^. Because double-stranded DNA prefers B-form geometry in solution, it must be converted to A-form upon binding in the central chamber. On the other hand, RNA-DNA hybrids are preconfigured as A-form. Thus, the clamp loader does not need to overcome this energetic penalty for loading at the RNA-primed p/t-junction. The only difference between loading complexes is that the RNA-primed p/t-junction does not move up as high in the central chamber of the clamp loader as a result of flexibility in the “separation pin” (**Fig. 3D, Fig. S7E**).

We then wondered whether these subtle differences in p/t-junction binding manifest as differences in clamp-loading competency or p/t-junction binding affinity. To test the ability to load clamps, we measured steady-state ATPase rates, which report on the turnover of the clamp loader upon clamp loading. A p/t-junction containing an RNA primer is slightly more effective at activating the clamp loader ATPase activity, although this effect disappears in the presence of ß-clamp (**Fig. 3E**). To test whether the primer composition affects binding affinity, we performed an anisotropy assay with TAMRA-labeled p/t-junctions. Surprisingly, we find that the affinity for an RNA-primed p/t-junction is identical to that of a DNA-primed junction (**Fig. 3F**). We speculate that the difference in registry of the p/t-junction within the central chamber provides additional binding energy for the DNA-primed substrate that offsets the energetic penalty of converting the DNA duplex to A-form.

#### Altered-Collar (overall resolution 3.8 Å)

A surprising class of DNA-primed particles exhibits an open clamp structure (**Fig. 4A**), but the collar region is in a different conformation from all other γ-complex structures reported herein or elsewhere^32,39,40^. The collar region of the B subunit shifts upward away from the AAA+ spiral. The collar regions of the C, D and E subunits are static, and the A subunit only moves slightly (**Fig. 4B**). The movement of the collar opens a large pore between the A and B subunits (**Fig. 4C**). A pore opens and closes at this site in the eukaryotic clamp loader RFC, although it manifests through a different mechanism^27–30^.

**Figure 4.**
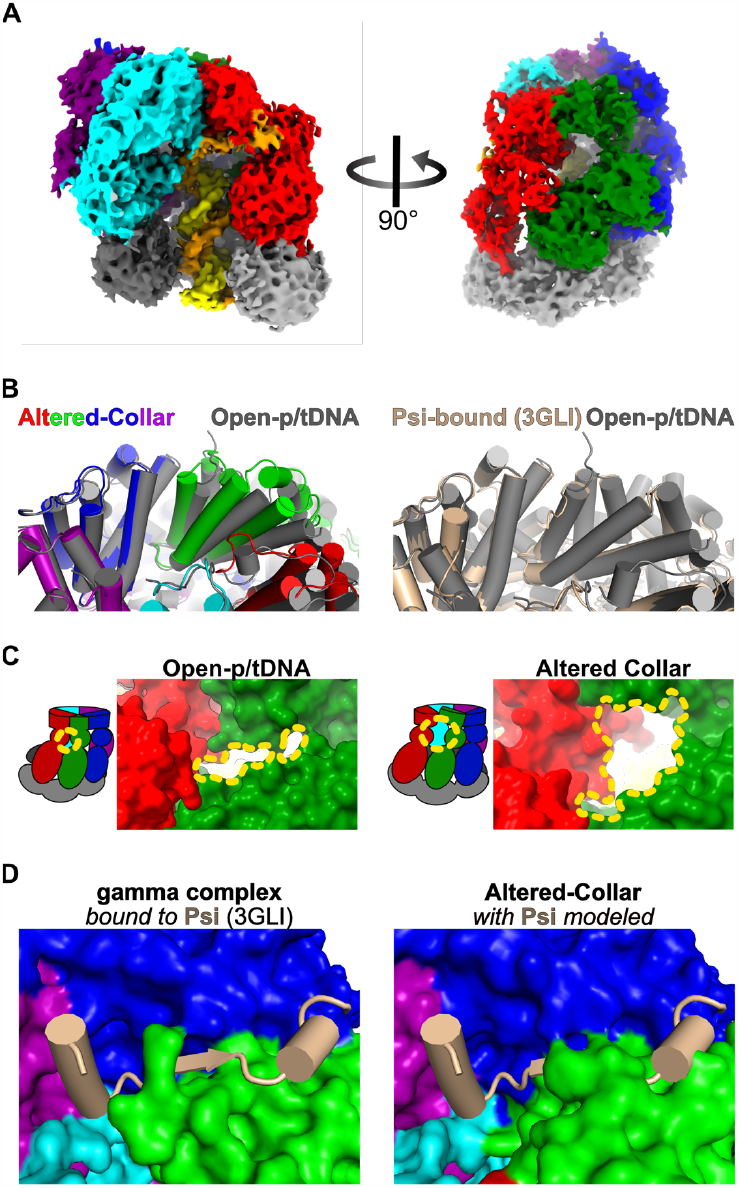
Structure of the Altered-Collar conformation of the bacterial clamp loader. **A)** Cryo-EM map of the Altered-Collar state (bound to p/t-DNA and ATP-analog). **B)** Comparison of collar conformation across states. All structures were aligned on the collar of the D subunit. *(Left)* The Altered-Collar (color) is compared to the Open-DNAp/t structure (gray), and *(Right)* Open-p/tDNA is compared to the psi-bound (wheat, PDB: 3GLI) state. **C)** The interface between A and B subunits. The Altered-Collar conformational change results in a large opening between the A and B subunits of the clamp loader (yellow dashed contour). **D)** Modeling the psi peptide into the Altered-Collar. *(Left)* A solved co-crystal structure of the clamp loader (surface) bound to psi peptide (cartoon). *(Right)* The co-crystal structure was aligned to the Altered-Collar structure as in panel B. This alignment was used to superimpose the psi peptide onto the Altered-Collar structure.

Moreover, this movement alters the center of the collar at the binding site for the ψ protein, an accessory factor that, together with the χ protein, acts as an adaptor linking the clamp loader to Single-Stranded Binding Protein (SSB)^39,41–43^. The Altered-Collar state shifts the binding site such that it is incompatible with the known binding pose of the ψ protein^39^ (**Fig. 4D**). We attribute this structure to an off- or parallel-pathway intermediate and discuss potential implications for the Altered-Collar conformational changes in the Discussion.

#### Closed-DNA1 & Closed-DNA2 (overall resolution 3.8 Å & 3.9 Å)

The Closed-DNA1 and Closed-DNA2 structures reveal how the clamp closes and yields insight into sliding clamp release (**Fig. 5A&B**). Both of these structures have a closed ß-clamp, which is in the same conformation as the free clamp (C_α_-RMSD 1.4 Å and 1.3 Å from PDB: 2POL). By comparing the Open-DNAp/t and Closed-DNA1 structures we infer that the clamp closes through two motions. The sliding clamp subunit bound to A, B, and C moves laterally, while the second sliding clamp subunit bound to C, D, and E pivots down and in towards the A-gate (**Fig. 5D**). In spite of this, the conformation of the clamp loader is essentially identical across Open-DNAp/t, Closed-DNA1, and Closed-DNA2 (**Fig 5C**). The only difference between Closed-DNA1 and Closed-DNA2 lies within the contacts between the clamp loader and the sliding clamp (**Fig. 5E&F**). In DNA-Closed1, all five subunits of the γ-complex contact the ß-clamp, although the D and E subunits lose substantial contact relative to the Open-DNAp/t state. The Closed-DNA2 state further loses contact between ß-clamp and subunits C, D, and E of the γ-complex (**Fig. 5F**). Between Closed-DNA1 and Closed-DNA2 the sliding clamp separates from the E subunit by ∼4 Å. We propose that the Closed-DNA1 conformation precedes the Closed-DNA2 conformation in a trajectory towards clamp release. The disengagement of the D and E subunits from the clamp is similar to that seen in RFC and PCNA^27^.

**Figure 5.**
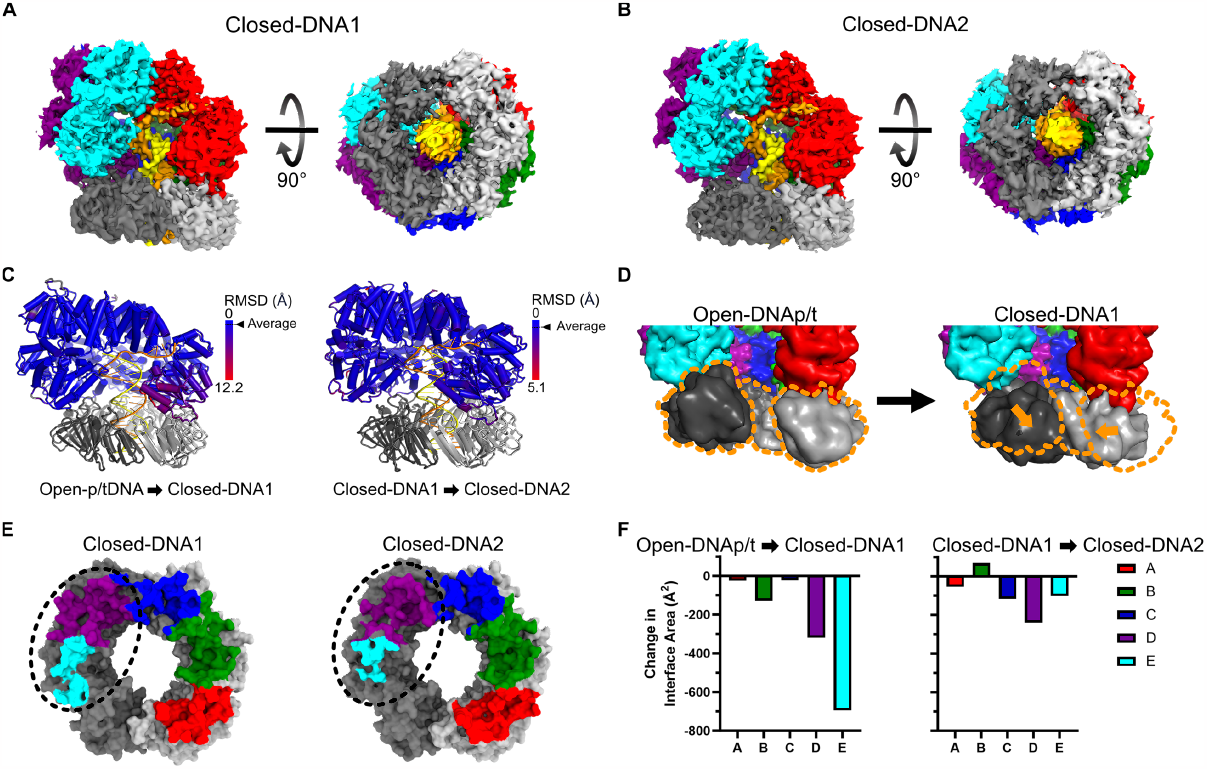
Closing the sliding clamp. **A-B)** Cryo-EM maps of the Closed-DNA1 and Closed-DNA2 conformations of the clamp loader and sliding clamp complex. **C)** Per-residue RMSD of the clamp loader comparing the Open-DNAp/t state to the Closed-DNA1 state and the Closed-DNA1 state to the Closed-DNA2 state. Structures were globally aligned and per-residue C_α_ RMSD was calculated. **D)** Comparison of the sliding clamp in open and closed conformations with p/t-DNA. Structures were aligned on the E subunit of the clamp loader and are shown as low contour surfaces. Center of mass shifts in the sliding clamp domain III of subunit I and domain I of subunit II are shown in orange vectors. The position of the clamp in the Open-DNAp/t conformation is outlined in orange to help guide the eye. **E)** Interface area between the sliding clamp and clamp loader. Models of the sliding clamp are shown as surfaces, and residues that contact the clamp loader are highlighted with color according to the contacting clamp loader subunit. The region that changes the most is highlighted with a dashed oval. **F)** The change in interface area between the clamp loader and sliding clamp during clamp closure.

In Closed-DNA1 and Closed-DNA2 structures, the clamp makes multiple direct contacts with DNA, primarily mediated by Gln15, Arg80 and Arg73 of clamp subunit I, and Gly23, Arg24, Arg80, and Gln149 of the second ß-clamp subunit. Previous work showed Arg24 and Gln149 of the ß-clamp are important for efficient loading^44^. Similar residues in PCNA contact DNA and have been shown to aid in clamp loading by RFC^27,45^. Our findings support their hypothesis that these residues contribute directly to the clamp closure step.

### Insights into the stimulation of ATPase activity

All structures reported here contain ATP analogs in the three ATPase active sites, which allows us to observe the steps that align the catalytic machinery for ATP hydrolysis. Prior to p/t-junction binding, the ATPase sites are not optimally positioned for ATP hydrolysis. In the Semi-Open state, there is a gap between the B and C subunits, which prevents the *trans*-acting arginine finger (Arg169 in γ, Arg158 in δ′) from interacting with ATP (**Fig. 2E**). In the Fully-Open state, the arginine finger of the C subunit contacts the B subunit’s ATP. Furthermore, we find no evidence for a previously hypothesized DNA-dependent allosteric switch involving Lys100 and Arg98 of γ acting in *cis*^*16*^, as this residue does not significantly alter its conformation upon DNA binding (**Fig. S8A**). Because DNA stimulates ATPase activity (**Fig. S1A**), we propose that p/t-junction binding aligns other residues that are necessary for ATP hydrolysis. We find that upon p/t-junction binding, a *trans*-acting lysine (Lys141 in γ, Lys130 in δ’) pivots upward to contact Asp126 of the Walker B motif in the neighboring active site (**Fig. S8B**).

We hypothesized that this *trans*-acting lysine plays a key role in stimulating ATPase activity in response to p/t-junction binding. To test this hypothesis, we made three variant clamp loader complexes in which this residue is mutated to alanine. The γ-K141A variant perturbs the active sites of the B and C subunits, while the δ’-K130A variant perturbs the D subunit’s active site. The γ-K141A/δ’-K130A double-mutant perturbs all three active sites. We find that all three variants have modestly impaired ATPase activity (**Fig. S8C**), and that addition of p/t-DNA boosts ATPase activity of all three variants in a manner similar to wild-type (**Fig. S8D**). These observations do not support our hypothesis that this lysine is the conformational trigger for ATPase activity.

## DISCUSSION

### Multi-step opening of the bacterial sliding clamp into a planar conformation

Our structures reveal how the *E. coli* clamp loader opens and places the ß-clamp onto DNA. By arranging our structures into a series of steps, we produce a nearly complete mechanistic description of the clamp loading reaction in bacteria (**Supplemental Video 1**). We divide the clamp loading cycle into five different phases i) clamp binding, ii) clamp opening, iii) p/t-junction binding, iv) clamp closure, and v) clamp loader recycling (**Fig. 6**). The structures reported here illuminate four of the five phases of the clamp loading cycle; future studies will focus on the clamp loader recycling phase.

**Figure 6.**
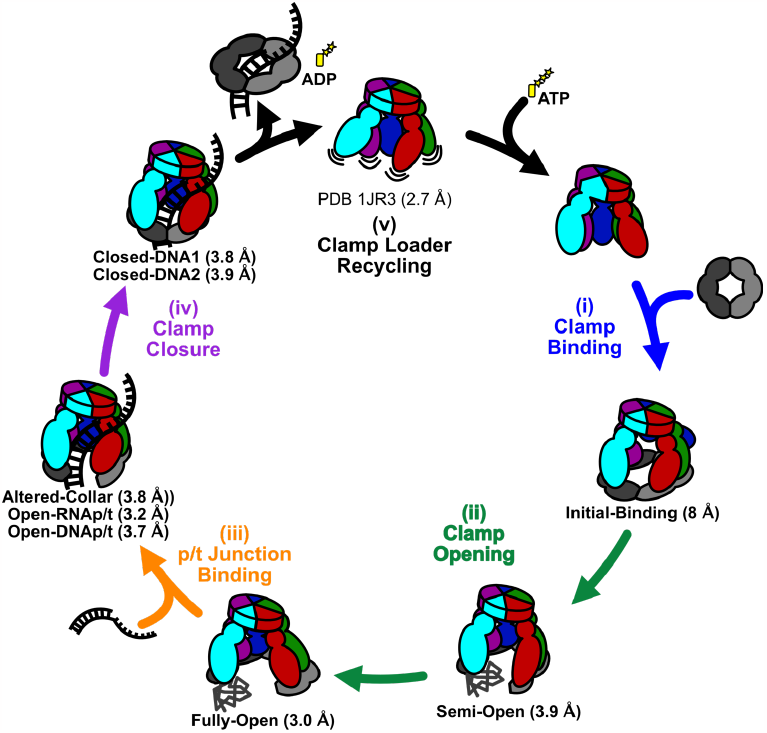
Updated bacterial sliding clamp loading mechanism. (i) Clamp loaders first bind ATP, then the sliding clamp. (ii) Sliding clamp opening occurs in two steps: first, a spiral state with a partial opening of the sliding clamp, and then a wide open state with a planar conformation. Domain I of subunit II of the sliding clamp is unstructured in both. (iii) The clamp loader/sliding clamp complex binds a p/t-junction and adopts a helical conformation. (iv) Binding of the p/t-junction triggers ATP hydrolysis and the sliding clamp closes around the p/t-junction. (v) The sliding clamp dissociates from the closed sliding clamp and ADP, and the loaded sliding clamp can be used by DNA replication and repair factors.

Our structure of the Initial-Binding state shows the clamp binding phase of the cycle. Despite the low resolution of the structure, we unambiguously observe that the γ-complex initially binds to a closed ß-clamp (**Fig. 2A; Fig. 6 i**). The existence of a closed binding complex was predicted based on rapid kinetics studies which showed that clamp binding occurs before clamp opening^46–49^. The γ-complex binds the ß-clamp using the A, D and E subunits, and with the A-gate of the γ-complex open. The open A-gate was predicted based on fluorescence resonance energy transfer (FRET) experiments^31^ and highlights one difference from the eukaryotic clamp loader, which binds to its clamp with a closed A-gate^27^.

The clamp opening phase (**Fig. 6 ii**) begins with the intermediate Semi-Open state, which was predicted by rapid-kinetics studies^50^. Here we observe that this intermediate has all five AAA+ modules of the γ-complex gripping the ß-clamp. The AAA+ modules adopt a spiral matching the tight spiral of the clamp. The clamp opening phase ends in the Fully-Open state, with the ß-clamp opened into a wide, planar arrangement (**Fig. 3B**). This conformational change is driven by a pivoting motion between the B and C subunits that flattens the spiral of both the AAA+ modules and ß-clamp as they open.

Our structures show that the clamp loader undergoes a dramatic conformational change when opening the sliding clamp. Paradoxically, a previous FRET study indicated that the distance between the A and E subunits does not change during opening, and it was interpreted that the γ-complex does not alter its conformation during clamp opening^31^. An equilibrium between the constricted Semi-Open and the splayed Fully-Open states could explain the FRET data. The inter-probe distance estimated from the FRET data is 50-52 Å. The estimated inter-probe distances in our Semi-and Fully-Open states are ∼40 Å and ∼55 Å, respectively. Taken together, the FRET data suggests that the Fully-Open state accounts for ∼75% of γ-complex bound to clamp in solution.

The ß-clamp lacks density for an entire domain at the clamp opening in the Semi- and Fully-Open states, suggesting that the domain unfolds. Unfolding of this domain upon ß-clamp opening was predicted from hydrogen-deuterium exchange data^36,37^. This unfolding changes the size of the opening in the clamp, potentially affecting how DNA enters into the central chamber. Unfolded proteins are more extended, so this domain may partially block the gap through which DNA must enter. We speculate that the wide opening in the Fully-Open state allows for p/t-junction passage despite the potential steric block of the unfolded domain.

The third phase of the clamp loading cycle (p/t binding) is illuminated by our structures of γ-complex and ß-clamp bound to p/t-junctions **(Fig. 6 iii)**. The p/t-bound state is nearly identical in the presence of either a DNA- or RNA-primer (**Fig. 3; Fig S7D**), providing confidence in experiments using DNA-primed p/t-junctions. After p/t-junction binding, the clamp loader AAA+ domains tighten around the DNA helix, constricting the whole complex. This constricted spiral state follows the DNA backbone and is distinct from that observed in the Semi-Open state (C_α_-RMSD ∼5.1 Å; **Fig S7C** ).

Binding of a p/t-junction stimulates ATPase activity^5^, but it remains unclear how. We previously proposed that specific residues would trigger ATP hydrolysis after a conformational change upon p/t-junction binding. We first predicted that γ Lys100 and γ Arg98 within the AAA+ module controls ATPase activity *in cis*^*3,16,18*^. However, we observed that γ Lys100 or γ Arg98 do not have the predicted conformational change between the Fully-Open and Open-p/t structures (**Fig. S8A**). On the other hand, *trans*-acting lysines (γ Lys141 and δ’ Lys130) directly contact the Walker B motif (Asp126) of the neighboring ATPase active site only after p/t-junction binding (**Fig. S8B**). These lysine residues have been proposed to play a role in ATP hydrolysis based on bioinformatic^51^ and deep mutational scanning analyses^52^. We find that clamp loader variants with these residues mutated are still able to efficiently stimulate ATPase activity in response to p/t-junction binding (**Fig. S8D**). This indicates that these lysines are not the DNA coupling element, as we hypothesized. However, the mutated variants have lowered overall ATPase activity (**Fig. S8C**), suggesting that this *trans*-interaction plays a role in supporting the active site. We also could not identify the trigger for DNA-dependent ATPase stimulation from similar structures of the eukaryotic clamp loader^27^.

The clamp closure phase of the cycle is illuminated by our structures of the γ-complex bound to a ß-clamp that is closed around p/t-DNA (Closed-DNA1 and Closed-DNA2) **(Fig. 6 iv)**. These structures show that clamp closure can occur with no significant structural change in the clamp loader. Nonetheless, there is a substantial loss of interaction of the D and E subunits with the sliding clamp. After clamp closure, the loader continues to disengage with the E, D and C subunits. We speculate that this is on-pathway towards complete disengagement of the clamp loader.

Moreover, these structures indicate that clamp closure does not require ATP hydrolysis. Prior experiments showed that ATP hydrolysis precedes clamp closure^14^. Our structures show that ATP hydrolysis is not strictly required for closure, but is more likely driving complete disengagement of the clamp loader. To reconcile our structures and the kinetic data, we note that all of the structures bound to p/t-junctions have their ATPase sites engaged, regardless of whether the p/t-junction contains an RNA or DNA primer, or whether the clamp is open or closed. Therefore, ATP hydrolysis presumably can manifest from any of these states, including prior to clamp closure.

### Different opening mechanisms between bacterial and eukaryotic clamp loaders

The *E. coli* clamp loader has been studied for decades and has been the archetype for bacterial clamp loaders^3,53^. Although eukaryotes have multiple clamp loaders^18,54^, their primary clamp loader is RFC, analogous to the γ-complex. Recent structural studies have revealed RFC’s mechanism in unprecedented detail^22,26–30^, which allows us to compare bacterial and eukaryotic clamp loaders.

The two clamp loaders bind to their sliding clamps differently during the clamp loading cycle, which may reflect differences within the sliding clamp itself (**Fig. 7A**). Each sliding clamp subunit has a strong partner binding site within one domain, while the remaining domains contain weak binding sites (**Fig. 7B**). These strong binding sites are used by other DNA replication and repair proteins. Because ß-clamp and PCNA have different stoichiometries (dimer vs. trimer), ß-clamp has two strong binding sites whereas PCNA has three. The different symmetries change the registry of the strong binding sites with the clamp loader subunits. For instance, ß-clamp’s strong binding sites are aligned with the A and D subunits of the clamp loader. Accordingly, γ-complex initially binds the ß-clamp primarily using its A, D and E subunits (**Fig. 7A**). In contrast, PCNA’s strong binding sites are aligned with the A, C, and E subunits of RFC, and RFC first binds to the clamp using its A and C subunits with contributions from the B subunit^22,26,27^.

**Figure 7.**
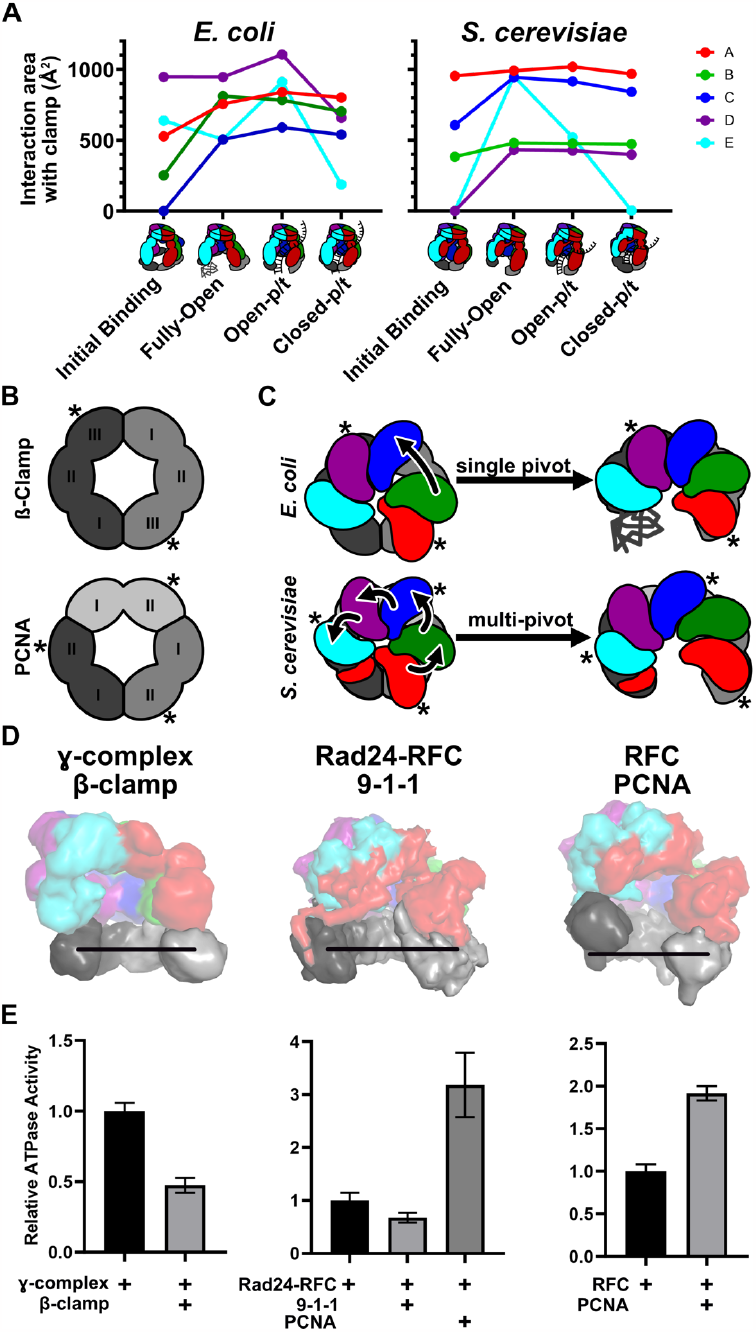
Comparing Bacterial and Eukaryotic clamp loader mechanisms. **A)** Interface area between the *E. coli* (left) and *S. cerevisiae* (right) clamp loader and sliding clamp at comparable stages during clamp loading^27^. When multiple structures correspond to one state, we report the average value across all structures. **B)** Location of strong binding domains of the sliding clamps are highlighted with asterisks. **C)** Comparison of the clamp opening motions of the bacterial clamp loader and RFC. The colored shapes represent the Rossmann domains of each subunit and the gray rings represent the sliding clamp (viewed from above). Arrows indicate the motion of the Rossmann domains transitioning from the initial binding to fully open conformations. **D)** Structures of the open bacterial clamp loader, Rad24-RFC (PDB: 7ST9), and RFC (PDB: 7TI8) complexes bound to their cognate sliding clamps (displayed as low contour surfaces). The solid line approximates the plane of the corresponding closed sliding clamp. **E)** The ATPase activity of the bacterial clamp loader, Rad24-RFC^56^, and RFC^7^, either alone or with a sliding clamp. ATPase activities are relative to the rate of the clamp loader alone.

In either case, clamp opening corresponds with complete engagement of the remaining clamp loader subunits with the sliding clamp. For the *E. coli* system, this means ß-clamp opening occurs primarily by engaging the B and C clamp loader subunits, whereas PCNA opening is concomitant with engagement of the D and E RFC subunits. Both loaders interact with the clamp most extensively in an open state. However, binding of a p/t-junction further increases the contacts between γ-complex and ß-clamp but causes RFC to lose interaction with PCNA. Upon clamp closure around p/t-junctions, both complexes lose substantial interactions between the sliding clamp and the E subunit. In general, the D subunit of γ-complex has the most extensive interaction with ß-clamp, while the A subunit of RFC has the tightest interaction with PCNA. This observation is surprising because the A subunit of the γ-complex (the δ protein) shows the tightest affinity with the sliding clamp^85^. We hypothesize that the different stoichiometry of the bacterial and eukaryotic sliding clamps and ultimately the positioning of strong binding sites is a major factor underlying the different binding and opening mechanisms.

The bacterial and eukaryotic sliding clamps also open into distinct conformations by distinct motions. The γ-complex opens its clamp in two steps, both of which have one domain of the sliding clamp unfolded. The clamp is first broken open with a downward motion into a spiral Semi-Open conformation. A large pivoting motion at a single interface between the B and C subunits transitions the Semi-Open conformation into the planar Fully-Open state (**Fig. 7C**). In contrast, RFC uses multiple pivoting motions between nearly all subunits to open PCNA into a spiral conformation with no obvious intermediates and no unfolding^27^. The planar opening of the bacterial ß-clamp is wider than that seen in the spiral PCNA (**Fig. 7D)**. In addition, the γ-complex’s lack of an A’ domain could allow for greater conformational flexibility leading to the wider A-gate opening seen in the *E. coli* clamp loader. We speculate that wider opening of the planar ß-clamp and A-gate allows for efficient DNA passage into the central chamber despite the steric block of the unfolded ß-clamp domain.

Our comparative study leads to a hypothesis for how the conformation of the open clamp alters ATPase activity. We hypothesize planar conformations inhibit ATPase activity because the active sites are misaligned. Conversely, a spiral configuration of AAA+ domains results in better alignment of the ATPase active sites^27^. In support of this hypothesis, addition of PCNA boosts RFC’s ATPase activity^7^, while addition of ß-clamp to γ-complex inhibits ATPase activity (**Fig. 7E &** ^5^). Moreover, Rad24-RFC decreases in ATPase activity in the presence of its cognate clamp (**Fig. 7D,E**&^56^), which it opens in a planar fashion^23,24^.

Are the different clamp opening mechanisms primarily driven by the energetics of the clamp loader or by that of the sliding clamp? We speculate that the sliding clamp plays the predominant role in dictating the opening geometry, although almost undoubtedly both components play some role. Molecular Dynamics (MD) simulations have suggested that PCNA prefers a right-handed helical conformation with multiple pivot points^57^, while the *E. coli* clamp prefers to adopt a planar conformation with pivoting at the same site as when bound to the γ-complex^58^. These same geometries are observed in structures of the open sliding clamps bound to their respective clamp loaders. Thus, the clamp loader does not appear to be imparting the conformation onto the sliding clamp. In further support of this hypothesis, the T4 bacteriophage sliding clamp (in the presence of p/t-DNA) is opened into a right-handed spiral by its cognate clamp loader^16^, and MD simulations show that the free T4 clamp prefers to adopt a right-handed spiral in the open form^59^. We also note that Rad24-RFC, which shares four out of five subunits with the RFC loader, opens the 9-1-1 clamp into a planar conformation^23,24,60^. Accordingly, we predict that the free 9-1-1 clamp prefers planar opening, even without the presence of the Rad24-RFC loader. Moreover, addition of PCNA to Rad24-RFC results in a boost in ATPase activity (**Fig. 7E**)^56^, presumably because PCNA prefers to open into a right-handed spiral and thus aligns the ATPase sites more optimally.

A final distinction between bacterial and eukaryotic clamp loaders is that RFC partially unwinds the p/t-junction^27^, while the *E. coli* loader has no obvious base-melting activity. In our structures or elsewhere^39^, we observe no unwinding of the p/t-junction, regardless if the primer is DNA or RNA. Moreover, our previous fluorescence data indicate that the *E. coli* clamp loader is unable to unwind a DNA p/t-junction^27^. RFC’s unwinding activity is thought to facilitate its role in loading at nicked DNA for performing long-patch Base Excision Repair^29,30,61^. Because the *E. coli* clamp loader is not known to play a role in Base Excision Repair, presumably it does not require DNA unwinding activity. Nonetheless, the *E. coli* γ-complex is competent to load ß-clamp onto nicked DNA^62^. This activity remains mysterious, as our structure suggests that at least five nucleotides of ssDNA in the template strand are necessary to exit the central chamber of the loader. We note that the *E. coli* clamp loader has flexibility in its separation pin element (**Fig. 3D**), which allows it to adopt two different heights of the primer 3’ end (**Fig. S7E**). We propose that this flexibility may play a role in loading at nicked DNA. Future studies will illuminate this mechanism.

### Speculation on the Role of the Alternate Collar Conformation

We observe a novel conformation of the collar region in the Altered-Collar class of particles that contains p/t-DNA. This conformational change is much larger than seen in previous structures, in which minor modification of the collar resulted in modulation of the ψ protein binding site^39^. The Altered-Collar state observed here severely disrupts the ψ protein binding site (**Fig. 4D**), suggesting that this conformation is incompatible with ψ protein binding. It is possible that the Altered-Collar state functions as a means for disengaging the clamp loader’s linkage to SSB through the ψ protein.

We also observe that a pore opens between the AAA+ modules of the A and B subunits (**Fig. 4C**). The position of this pore is similar to a pore that we recently discovered in the yeast RFC complex that is important for loading onto nicked DNA for repair^27–30^. Because the *E. coli* clamp loader can readily load ß-clamp onto nicked DNA^62^, we speculate that this pore could function as a channel for the unwound primer strand of nicked DNA to exit the central chamber, similar to RFC. Future experiments will test these two non-mutually exclusive hypotheses.

### Conclusions

Altogether, our comparative study of clamp loading in bacteria and eukaryotes reveals the intricate molecular choreography that underlies clamp loading over one billion years of evolution. We find numerous distinctions between bacterial and eukaryotic clamp loading, despite the universal conservation of clamp loading throughout life. Our work opens the door for potentially targeting the clamp loading process of bacteria for development of novel antibiotics.

## Supporting information

Initial-Binding

Semi-Open

Fully-Open

Open-DNApt

Open-RNApt

Closed-DNA1

Closed-DNA2

Altered-Collar

## AUTHORS’ NOTE

During the preparation of this manuscript, the authors became aware of a preprint^63^ that describes similar structures as those we report here. In order to not bias our interpretation of our data, we did not read that paper in detail until after our manuscript was in its final form. We note that many of our structures broadly match those described in the preprint^63^.

## MATERIALS AND METHODS

### Expression Plasmid Cloning

#### δ pET28PP

All primers used for cloning are listed in Supplemental Table 1. To make the expression plasmid for δ protein, we transferred the δ coding region from pET3c^39^ to pET28PP for higher expression using standard restriction enzyme cloning. We generated the δ coding region with flanking cleavage sites for the restriction enzymes Xbal and BamHI using PCR with Q5® High-Fidelity DNA Polymerase and DNA oligonucleotides (Integrated DNA Technologies). PCR products were purified with the Monarch PCR&DNA Cleanup kit (New England Biolab). The pET28 vector was purified from DH5a *E. coli* (Thermo Fisher Scientific) using a Qiagen miniprep kit (Thermo Fisher Scientific). Both the δ insert and pET28 plasmid were digested with the restriction enzymes XbaI and BamHI (New England Biolabs) and then purified by gel extraction using the QIAquick Gel Extraction kit (Qiagen). The δ coding region was ligated into pET28PP using T4 DNA ligase in T4 DNA ligase buffer (New England Biolabs). This mixture was transformed into Mix & Go competent DH5a *E. coli* (Zymo Research) and plasmids were recovered by miniprep (Qiagen). Plasmids were checked for proper gene insertion by Sanger sequencing (Genewiz).

#### His-tagged δ’ pET28PP

Gibson assembly was used to generate a 6x-His tag δ’ pET28PP plasmid for expression of the δ’ protein.

The pET28 backbone was generated by PCR. The δ’ insert was amplified by PCR from a δ’ containing pET3a vector. After purification of the PCR products, the δ’ fragment was inserted into the pET28PP backbone via Gibson assembly (2x NEBuilder Hifi DNA Assembly Mix, New England Biolabs). Expression plasmids were transformed into Mix & Go competent DH5a E. coli and plasmids were recovered by miniprep (Qiagen). Plasmids were checked by Sanger sequencing (Genewiz).

#### δ’-K130A and γ-K141A pET28PP

Inserts with the coding region for δ’-K130A and γ-K141A and flanking regions containing restriction digest sites, were purchased from Twist Bioscience. We treated the inserts and the pET28PP backbone with the restriction enzymes XbaI and XhoI and then purified the products by gel extraction using the QIAquick Gel Extraction kit (Qiagen). The coding regions were ligated into pET28PP using T4 DNA ligase in T4 DNA ligase buffer (New England Biolabs). These mixtures were transformed into chemically competent 10-β *E. coli* (New England Biolabs) and plasmids were recovered by miniprep (Qiagen). Plasmids were checked for proper gene insertion by Sanger sequencing (Genewiz).

### Expression and Purification of *E. coli* Clamp Loader

The γ, γ-K141A, δ, δ’, and δ’-K130A proteins of the clamp loader and the β-clamp were expressed individually in the BLR21(DE3) strain of E. *coli*. Plasmids for expression of the β and γ proteins are as described in ^39^. We used a truncated form of the γ subunit (residues 1-373), which removes the segments that bind to the polymerase and helicase^64,65^ and are dispensable for clamp loading^66^. The γ, δ’, γ-K141A, δ’-K130A, and β proteins each contain a Prescission Protease cleavable 6x-His tag at their N-terminus.

#### Protein Expression

The expression plasmids were separately transformed into BLR21(DE3) *E. coli* cells (Millipore) and grown overnight at 37 °C and then transferred into 100 ml Luria Broth media supplemented with 100 μg/ml Kanamycin pre-warmed to 37 °C. These cultures were grown for 4 hours while shaking at 37 °C. Then 20 mL of the starter cultures were used to inoculate 1 L of Terrific Broth media pre-warmed to 37 °C. The cultures were grown until they had an optical density at 600 nm of 0.6-0.8 and then were induced with isopropyl β-D-1-thiogalactopyranoside (IPTG) at 1 mM final concentration. Following induction, the cultures were grown overnight at 18 °C. Cells were then harvested by centrifugation at 5,000 x g for 30 minutes. Cell pellets were flash frozen in liquid nitrogen and then stored at -80°C.

##### Protein purification

To purify the His-tagged proteins (γ, δ’, and β), the frozen pellets were thawed on ice and then resuspended in 20 mM Tris pH 8.0, 500 mM NaCl, 10% glycerol (v/v), 5 mM β-mercaptoethanol (BME), and 20 mM imidazole (Buffer Ni-A) and lysed with a cell homogenizer (Microfluidics Inc, Westwood, MA). Cell lysates were then clarified by centrifugation at 20,000 x g for 20 minutes. Lysates were then loaded onto HisTrap columns (Cytiva) pre-equilibrated with Buffer Ni-A. The loaded HisTrap columns were washed with at least 5 column volumes of Buffer Ni-A. The protein was then eluted with 20 mM Tris pH 8.0, 500 mM NaCl, 10% glycerol (v/v), 5 mM BME, and 250 mM imidazole (Buffer Ni-B). Fractions were then checked for protein by SDS-PAGE and pooled.

#### δ’ protein

The His-tag on δ’ was left intact. His-δ’ was dialyzed into buffer Ni-A overnight, then concentrated to ∼15 mg/ml, flash frozen, and stored at -80 °C.

#### γ protein

His-γ was treated with PreScission Protease to remove the His-tag, while dialyzing into Buffer Ni-A overnight at 4 °C. The treated protein was then passed over a HisTrap column, and the flowthrough was collected. Flowthrough fractions containing purified γ were pooled and dialyzed into 50 mM Tris pH 7.5, 50 mM NaCl, 10% glycerol, and 2 mM dithiothreitol (DTT) (Storage Buffer) overnight at 4 °C. Finally, γ protein was concentrated to ∼6 mg/ml, flash frozen, and stored at -80 °C.

#### β protein

Fractions containing His-β protein were pooled and dialyzed into 50 mM Tris pH 8.0, 10% glycerol, 2mM DTT overnight at 4°C while being treated with Prescission Protease. The protein was then loaded onto a 5-ml HiTrap Q column (Cytiva). A gradient of 50 mM Tris pH 8.0, 500 mM NaCl, 10% glycerol, 2mM DTT was used to elute β. Fractions containing β were pooled and dialyzed into Storage Buffer overnight. Purified β protein was concentrated to ∼16 mg/ml, flash frozen, and stored at -80 °C.

#### γ-K141A and δ’-K130A

To purify the His-tagged γ-K141A and δ’-K130A mutants, the frozen pellets were thawed on ice and then resuspended in Buffer Ni-A, and then lysed with a LM10 Microfluidizer^®^ Processor (Microfluidics). Cell lysates were then clarified by centrifugation at 20,000 x g for 20 minutes. 10 ml of Nickel NTA Agarose Beads (Gold Biotechnology) was added to each clarified lysate. The nickel resins were incubated with the supernatants for 1 hour at 4 °C while mixing. The resins were then collected by centrifugation and washed with at least 5 CV of Buffer Ni-A. 10 ml of Buffer Ni-B was added to the resins and the resins were incubated for 30 minutes at 4 °C while constantly mixing. After the incubation, the supernatants were collected and checked for protein by SDS-PAGE. Both the γ-K141A and δ’-K130A supernatants contained protein and the proteins were dialyzed into Buffer Ni-A overnight while being treated with Prescission Protease.

After Prescission Protease treatment, the proteins were collected and loaded onto HisTrap columns (Cytiva) pre-equilibrated with Buffer Ni-A. A 0-100% gradient of Ni-B was applied to the columns. γ-K141A and δ’-K130A eluted between 7-20% and 20-40% Buffer Ni-B, respectively. To check for cleavage of the His-tag, protein fractions were assessed by SDS-PAGE and near complete cleavage of the tag was observed. The purified γ-K141A and δ’-K130A were then dialyzed into 50 mM Tris pH 7.5, 50 mM NaCl, 10% glycerol (v/v), and 2 mM DTT (Buffer Q-A) at 4 °C overnight. Following dialysis, γ-K141A and δ’-K130A were concentrated to ∼5 and ∼2 mg/ml, respectively, flash frozen, and stored at -80 °C.

#### δ protein

To purify δ protein, the frozen cell pellets were resuspended in 20 mM Tris pH 7.5, 10% glycerol, 0.5 mM Ethylenediaminetetraacetic acid (EDTA), and 2 mM DTT and lysed using a cell homogenizer (Microfluidics Inc, Westwood, MA). Cell lysate was clarified by centrifugation at 20,000 x g for 20 minutes. The clarified lysate was then loaded onto a 5 ml HiTrap SP-sepharose column (Cytiva). δ protein was eluted with a gradient of 20 mM Tris pH 7.5, 500 mM NaCl, 10% glycerol (v/v), 0.5 mM EDTA, and 2 mM DTT over a 10 column volume gradient. Fractions containing δ protein were pooled and concentrated. Purified δ protein was then dialyzed overnight into 50 mM Tris pH 7.5, 300 mM NaCl, 10 % glycerol (v/v), 2 mM DTT, and 0.5 mM EDTA. Following dialysis, the protein was concentrated to ∼2.0 mg/ml, flash frozen, and stored at -80 °C.

#### γ complex

To reconstitute the γ-complex, we combined purified His-δ’, δ, and γ proteins in a stoichiometric ratio of 0.75:1:3, and the protein mixture was then dialyzed overnight into Buffer Ni-A. We then loaded the protein onto a 5 mL HisTrap column (Cytiva), then eluted with Buffer Ni-B. Fractions containing the assembled complex were combined with Prescission Protease, and were dialyzed overnight into Buffer Ni-A. The γ-complex was then passed over a 5 mL HisTrap column, and the flowthrough was collected and dialyzed into Buffer Q-A. The dialyzed protein was then loaded onto a 5 mL Q-sepharose column (Cytiva) and eluted with a 10 column volume gradient of 50 mM Tris pH 7.5, 500 mM NaCl, 10% glycerol (v/v), and 2 mM DTT (Buffer Q-B). Fractions containing the intact γ-complex were then pooled and dialyzed into Storage Buffer. We then concentrated the purified γ-complex to ∼2.5 mg/ml using an Amicon Ultra-15 centrifugal filter with a 30,000 kDa molecular weight cutoff. Aliquots were flash frozen in liquid nitrogen and stored at -80 °C.

#### γ-K141A and δ’-K130A γ complexes

To study the effect of the γ-K141A and δ’-K130A mutations on the γ complex, we purified WT γδδ’, γ_K141A_δδ’, γδδ’_K130A_, and γ_K141A_δδ’_K130A_ complexes. The γ_WT/K141A_, δ, and δ’_WT/K130A_ proteins were combined in a 2.5:1:1 stiocheometric ratio and dialyzed into Q-A overnight at 4 °C. The next day, the protein was loaded onto a 1 ml Hitrap Q column (Cytiva). The column was washed with at least 5 column volumes of Buffer Q-A, and then, the protein was eluted with a 10 column volume gradient of Buffer Q-B. Fractions containing the intact γ-complex were then pooled and concentrated. Aliquots of the purified complex were flash frozen in liquid nitrogen, and stored at -80 °C.

### DNA/RNA oligonucleotide preparation

DNA and RNA oligonucleotide sequences are listed in Supplemental Table 1. The RNA primer sequence was the preferred length and sequence of the *E. coli* primase^67^. Oligonucleotides were resuspended to a concentration of 100 μM in nuclease free water. To assemble the p/t substrates, the oligonucleotides were combined at equimolar concentrations (final stock concentration of 20-30 μM) in 25 mM Tris pH 7.5 and 5 mM MgCl_2_. To promote proper annealing, the equimolar oligonucleotide solutions were heated to 95°C for 2 minutes, then 85 °C for 3 min, then 75 °C for 5 minutes, then 65 °C for 5 minutes, then 55 °C for 10 minutes, then 50 °C for 10 minutes, then 45 °C for 10 minutes, then 40 °C for 10 minutes, then 35 °C for 10 minutes, then 30°C for 10 minutes, and 20 °C in a BioRad T100 Thermal Cycler until they were removed and stored at -20 °C.

### Cryo-EM sample preparation

For all samples, we first combined clamp loader and sliding clamp. Next, we added the p/t-junction, if present. We then added the ATP analog, which is either ATPγS or ADP•BeF_x_. If the sample is cross-linked, we then treat the sample with 1 mM of bis(sulfosuccinimidyl)suberate (BS3, Thermo Scientific Pierce) for 15 minutes at room temperature. To neutralize the cross-linking reaction, the samples were treated with Tris-HCl to a final concentration of 25 mM. Samples were stored on ice until grid preparation.

#### Clamp loader, sliding clamp, ATPγS sample

We combined equimolar clamp loader and sliding clamp to a final concentration of 3 μM each, then ATPγS was added to a final concentration of 1 mM. The sample was crosslinked according to the method above.

#### Clamp loader, sliding clamp, ADP•BeF_x_ sample

We made a solution 2.5 μM of clamp loader and 2.2 μM of sliding clamp. 1 mM of ADP•BeF_x_ was prepared in the sample according to (Kelch et al. 2011)^16^. The sample was then crosslinked according to the method above.

#### Clamp loader, sliding clamp, RNA/DNA p/t-junction, ADP•BeF_x_ sample

The sample was prepared such that the final concentration was 2.5 μM of clamp loader, 2.2 μM of sliding clamp, and 7 μM of either RNA/DNA p/t substrate or DNA/DNA p/t substrate. The oligonucleotide sequences are listed in Supplemental Table 1. 1 mM of ADP•BeF_x_ was prepared as described above. The RNAp/t sample was then crosslinked according to the method above.

### Cryo-EM grid preparation

All grids were first washed with ethyl acetate. The grids were then glow discharged on a PELCO easiGlow for 20-45 s at 10-25 mA (Ted Pella) (negative polarity). 3.5-4.5 μL of sample was applied to a grid and the grid was then blotted on both sides with a blot force of 5, a blot time between 5-7 seconds, and wait time between 0-2 seconds. The grids were then vitrified by plunging into liquid ethane using a Vitrobot Mark IV (FEI) at 10 °C and 95% humidity. For the specific grid type, glow discharge conditions, sample volume, blot force, and blot time for each sample see Supplemental Table 2.

### Cryo-EM data collection

#### Clamp loader, sliding clamp, and ATPγS sample

This sample was imaged on a Talos Arctica operated at 200 kV equipped with an GIF energy filter at x45000 magnification and a pixel size of 0.435 Å (bin=0.5) using a K3 Summit direct electron detector (Gatan) in superresolution counting mode. 3,664 micrographs were collected at a target defocus range of -1.1 to -2.3 and a total exposure dose of ∼43 e^-^/Å^2^ averaging 30 frames.

#### Clamp loader, sliding clamp, and ADP•BeF_x_ sample

This sample was imaged on a Titan Krios operated at 300 kV equipped with an GIF energy filter at x105000 magnification and a pixel size of 0.415 Å (bin=0.5), using a K3 Summit detector in superresolution counting mode. Movies were collected using SerialEM^68^. 4791 micrographs were collected at a target defocus range of -1.0 to - 2.4 and a total exposure dose of 49-50 e^-^/Å^2^ averaging 30 frames.

#### Clamp loader, sliding clamp, p/t-DNA, ADP•BeF_x_ sample

This sample was imaged on a Talos Arctica operated at 200 kV equipped with a Gatan K2 Summit direct electron detector and a Gatan energy filter. Micrographs were collected at x45000 magnification and a pixel size of 0.435 (bin=0.5), using a K3 detector in super-resolution mode. Movies were collected using SerialEM^68^. 3,846 micrographs were collected at a target defocus range of -1.0 to -2.4 and a total exposure dose of 49-50 e^-^/Å^2^ averaging 28 frames.

#### Clamp loader, sliding clamp, p/t-RNA, ADP•BeF_x_

This sample was imaged on a Glacios operated at 200 kV equipped with a Falcon4 (Thermo Fisher) direct electron detector and a Selectris energy filter operated at 10 eV. The Falcon4 camera was operated in electron event representation (EER) mode^69^. Micrographs were collected at x130000 magnification and a pixel size of 0.88.

Movies were collected using SerialEM^68^. 3,132 micrographs were collected at a target defocus range of -1.1 to -2.3 and a total exposure dose of ∼45 e^-^/Å^2^ averaging 1975 frames.

### Cryo-EM data processing

#### Clamp loader, sliding clamp, ATPγS

Micrographs were aligned in IMOD with 2x binning, resulting in a pixel size of 0.87 Å per pixel. After micrograph alignment, all further data processing was completed within cryoSPARC^70^. The initial Contrast Transfer Function (CTF) estimation was done using CTFFind4^71^. Following CTF correction, one round of Blob picking (90-160 Å) was performed to complete reference free particle picking. Particles were then extracted with a box size of 256 pixels. To identify quality particles, we performed two rounds of 2D classification and selected classes with well-defined features, resulting in a stack of ∼75,000 particles. This particle stack was used to train a Topaz particle picking model^72,73^. We implemented this Topaz particle picking model and identified 424,836 particles, which were extracted (box size 300 pixels). The particles were then subjected to 2 rounds of 2D classification and class selection. 2D classes with well-defined features were selected and then used to generate three *ab initio* models. The highest quality *ab initio* model appeared to be of a clamp loader/sliding clamp complex, and its associated particle stack was selected for further processing (211,589 particles). We then 3D classified this particle stack. We performed homogeneous refinement on the final particle stack of 44,484 particles. The B-factor used was 622.7 and the cryo-EM map had a resolution of 7.7 (FSC=0.143). All reference models used during data processing were downfiltered to at least 30 Å. A data processing pipeline is provided in Fig. S2 and data collection/3D volume reconstruction details are provided in Supplemental Table 3.

#### Clamp loader, sliding clamp, ADP•BeF_x_

Cryo-EM data processing was performed within cryoSPARC^70^. Cryo-EM movies were patch motion corrected followed by CTF correction using cryoSPARC’s implementation of Patch CTF. Blob picker was used for the initial particle picking (90-150 Å), which was followed by particle extraction (box size=256 pixels). These particles were then 2D classified and 2D classes with well-defined features were selected. These particles were then used as templates in the “Template Picker” tool and identified particles were extracted (box size=352). Another round of 2D classification followed by 2D subset selection was performed on the new particles. After 2D classification, the particle stack was used to generate 4 *ab initio* models. One of the *ab initio* classes was of a clamp loader/sliding clamp complex. This particle stack was then subjected to several rounds of 3D classification and homogeneous refinement to identify different conformations of the complex. A focus-mask was applied to the sliding clamp during the initial 3D classification to identify any open and closed states. This was followed by sequential non-masked 3-D classification. Non-uniform refinement was used for the 3D refinement of the final map^74^. A data processing pipeline is provided in Fig. S4 and data collection/3D volume reconstruction details are provided in Supplemental Table 3.

#### Clamp loader, sliding clamp, ADP•BeF_x_, and p/t-DNA

Cryo-EM data processing was completed within cryoSPARC^70^. Cryo-EM movies were patch motion corrected followed by CTF correction using cryoSPARC’s implementation of Patch CTF. Blob picker was used for the initial particle picking (90-150 Å), which was followed by particle extraction (box size=256 pixels). These particles were then 2D classified and 2D classes with well-defined features were selected. Particle picking was then performed using these particles as templates in the “Template Picker” tool and identified particles were then extracted (box size=300). Multiple rounds of 2D classification and 2D subset selection were performed to identify high-quality particles. After 2D classification, the particle stacks were used to generate 4 *ab initio* models. One of the *ab initio* models was of a clamp loader/sliding clamp complex. To identify the Open-DNAp/t, Closed-DNA1, and Closed-DNA2 states, 3D classification was first performed by applying a mask over the sliding clamp and A and B subunits of the *ab-initio* reconstruction. Subsequent 3D classification was performed without applying a focus-mask. To identify the Altered-Collar state, 3D classification was first performed by applying a mask over the collar of the *ab-initio* reconstruction. Non-uniform refinement was used for the refinement of the final maps^74^. A data processing pipeline is provided in Fig. S6 and data collection/3D volume reconstruction details are provided in Supplemental Table 3.

#### Clamp loader, sliding clamp, ADP•BeF_x_, and p/t-RNA

Cryo-EM data processing was completed within cryoSPARC^70^. Cryo-EM movies were patch motion corrected followed by CTF correction using cryoSPARC’s implementation of Patch CTF. Blob picker was used for the initial particle picking (90-150 Å), which was followed by particle (box size=260 pixels). These particles were then 2D classified and 2D classes with well-defined features were selected. Particle picking was then performed using these particles as templates in the “Template Picker” tool and identified particles were extracted (box size=260 pixels). These particles were then 2D classified, which was followed by 2D subset selection. After 2D class selection, the particles were extracted (box size=300 pixels, binning 2x). The resulting particle stack was used to generate 4 *ab initio* models. One of the *ab initio* models was of the clamp loader, sliding clamp, p/t-RNA complex. This particle stack was extracted (box size=300 pixels, binning=1x). The fully extracted particles were then 3D classified without a reference volume into 5 classes. No major differences were observed between the best three classes and they were pooled and refined by non-uniform refinement^74^. The map was sharpened with a B-factor of -92.3 within cryoSPARC and has a resolution of 3.2 Å (FSC=0.143). A data processing pipeline is provided in Fig. S5 and data collection/3D volume reconstruction details are provided in Supplemental Table 3.

### Model Building, refinement, and validation

We used crystal structures of the γ-complex bound to DNA-p/t (3GLF)^39^ and the β-clamp (2POL)^1^ as initial models. Each subunit was split into globular domains and then each domain was fit into the cryo-EM maps using UCSF ChimeraX^75^. Molecular Dynamics flexible fitting (MDFF) as implemented in NAMD 2.14 was used to generate the model of the Initial-Binding complex from the clamp loader, sliding clamp, ATPγS dataset^34,76^. Input files were prepared *via* the MDFF plugin in VMD 1.9.4a55^77^. The rigid body fit model and cryo-EM map were used as the starting model and bias potential. The protein interactions were described with the CHARMM36 force field^78^, and additional restraints added to maintain secondary structure, prevent *cis* peptide bonds, as well as maintain chirality. All other models were built using a combination of Coot for manual building^79^, Starmap for automated building^80^, and Phenix real-space refinement^81^. After the initial rigid body fitting, the Semi-open, Fully-Open, Open-DNAp/t, Open-RNAp/t, Closed-DNA1, and Altered-Collar models were adjusted in Coot. Following manual adjustment, the models and maps were used as inputs for Starmap, for rosetta-driven molecular structure refinement. The models from the final iterations of Starmap, were inspected and manually adjusted in Coot. The Closed-DNA1 Starmap output model was fit into the Closed-DNA2 map, the fitted model and map were then used as inputs for Starmap. The model from the final Starmap iteration for the Closed-DNA2 model was then manually adjusted in Coot. After all models were inspected and adjusted in Coot, they were refined using Phenix real-space refine. Errors in the models after refinement were manually fixed in Coot, and then a final round of real-space refinement was performed on each model. Real-space refinement statistics are listed in Supplemental Table 3. Figures and analyses were made using Pymol^82^, UCSF ChimeraX, and the PISA server^83^.

### ATPase assays

0.3 μM γ-complex was incubated with 0.2 μM (**Fig. S1B**) or 1 μM (**Fig. 3E, Fig. 7E, Fig. S8C**) ß-clamp, 0.4 μM (**Fig. S1B, Fig. 3E, Fig. 7E**) or 1 μM (**Fig. S8C**) p/t-junction and a mix of 5 U/ml Pyruvate kinase, 5 U/ml lactate dehydrogenase, 1 mM ATP, 150 μM phosphoenol pyruvate, 50 μM NADH, 25 mM Tris (pH 7.5), 100 mM potassium glutamate, 5 mM magnesium chloride. The oligonucleotide sequences are listed in Supplemental Table 1. The ATPase rates were measured at 25 °C with a VICTOR Nivo multimode plate reader (Perkin Elmer) to detect NAD+. Rates were obtained from a linear fit of the data in GraphPad Prism.

### Fluorescence measurements

To measure the affinity of different p/t-junction substrates to the clamp loader, sliding clamp, ADP•BeF_x_ complex we used a fluorescence anisotropy-based binding assay^84^. We used 5’ TAMRA-labeled template DNA which was annealed to either an 11 bp DNA or RNA primer, similar as previously described^16^. The oligonucleotide sequences are listed in Supplemental Table 1. Anisotropy measurements were made with a FluoroMax 4 (Horiba Jobin Yvon Inc) at 25 °C. Fluorescence was excited with a wavelength of 550nm (2 mm slit width) and emission was detected at a wavelength of 580 nm (7 mm slit width). Buffer conditions were 50 mM Tris (pH 7.5), 250 mM potassium glutamate, 2 mM DTT, 10% glycerol, and 1 mM magnesium chloride. 1 mM of ADP•BeF_x_ was prepared in the sample as described above. Then 5 μM sliding clamp and 150 nM p/t-junction substrate was added. Finally, the clamp loader was titrated into the solution. Raw data were fit using Prism (GraphPad Software) with the equation:

Y = Bmax*Xh / (Kdh + Xh) + Z, where X is the clamp loader concentration, Y is the anisotropy, h is the Hill slope, Z is the starting anisotropy, and Bmax is the maximum anisotropy.

## ACKNOWLEDGMENTS

The authors thank Drs. C. Xu, K. Song, K. Lee, and C. Ouch of the UMass Chan Medical School cryo-EM core facility for assistance with data collection. MDFF simulations were performed on the UMass Scientific Computing for Innovation Cluster. We thank Drs. N. Grigorieff, Y. Liu and members of the Kelch, and Schiffer Labs for helpful discussions. This work was funded by American Cancer Society grant PF-22-114-01-DMC to J. Pajak, and NIGMS grants R01-GM127776 and NIGMS R01-GM145943 to B. Kelch.

## AUTHOR CONTRIBUTIONS

J.L. and B.A.K. conceptualized research. J.L. performed all of the experiments with some assistance from E.K.N. and J.P.. J.L., J.P. and B.A.K. analyzed the data. J.L., J.P., E.L.S., and B.A.K. wrote the manuscript.

---

**Supplemental Figure 1.**
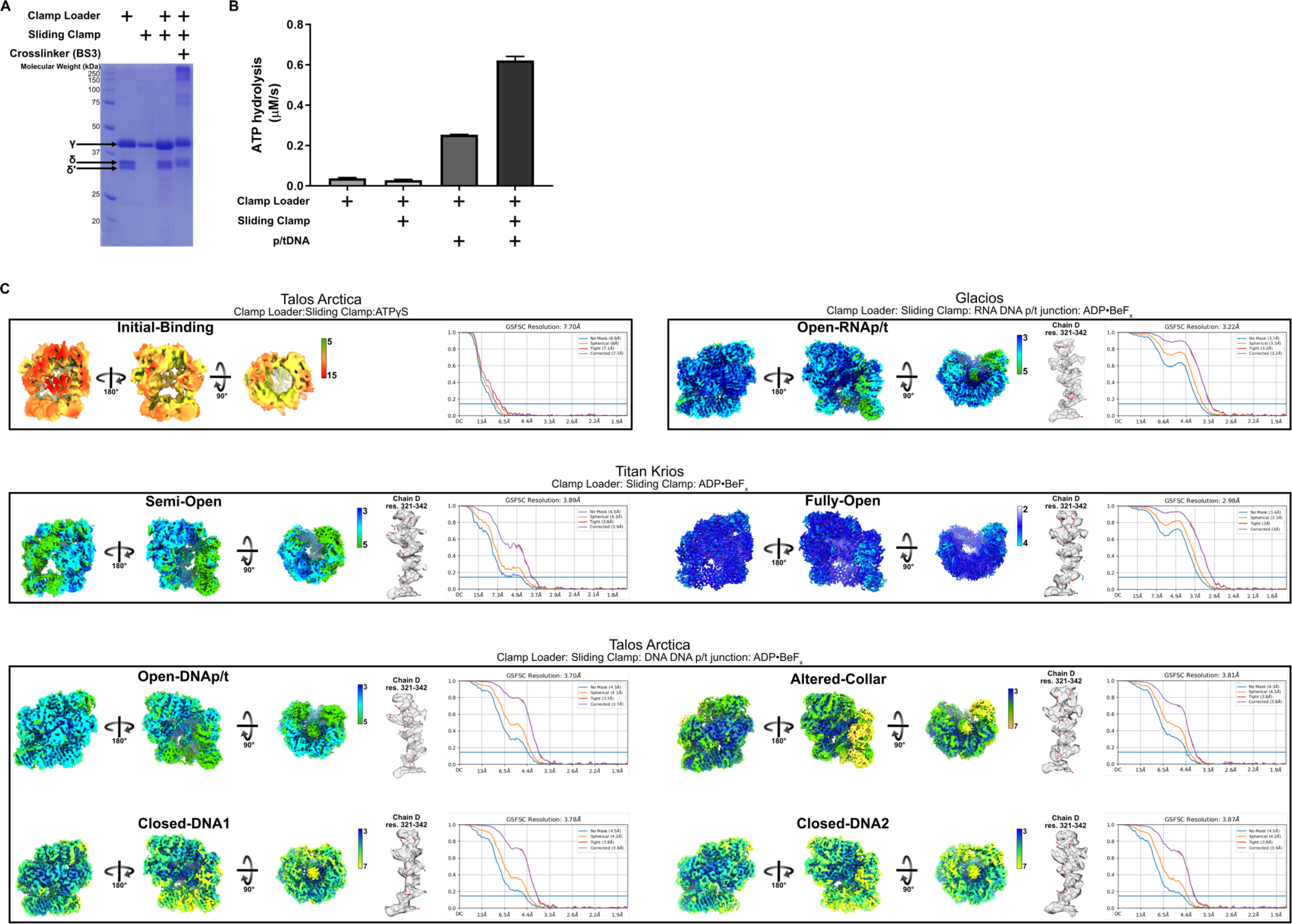
Characterization and cryo-EM map validation of the bacterial clamp loader. **A)** *Purified γ complex and β-clamp*. Sodium dodecyl sulfate-polyacrylamide gel electrophoresis (SDS-PAGE) gel of the purified γ-complex and β-clamp proteins, and samples of the γ complex combined with the β-clamp before and after crosslinking with BS3 crosslinker. **B)** *ATP activity profile of the bacterial clamp loader*. Steady-State ATPase activity of the purified clamp loader complex in combination with the sliding clamp and p/t-DNA. **C)** *Local resolution and FSC curves of the clamp loader cryo-EM maps*. Local resolution (Fourier shell correlation (FSC)=0.5) of the reconstructions and a representative section of each density map with fitted model. The overall resolution of each map was determined by the FSC of each half-map using Gold-standard cutoff of 0.143 (blue line).

**Supplemental Figure 2.**
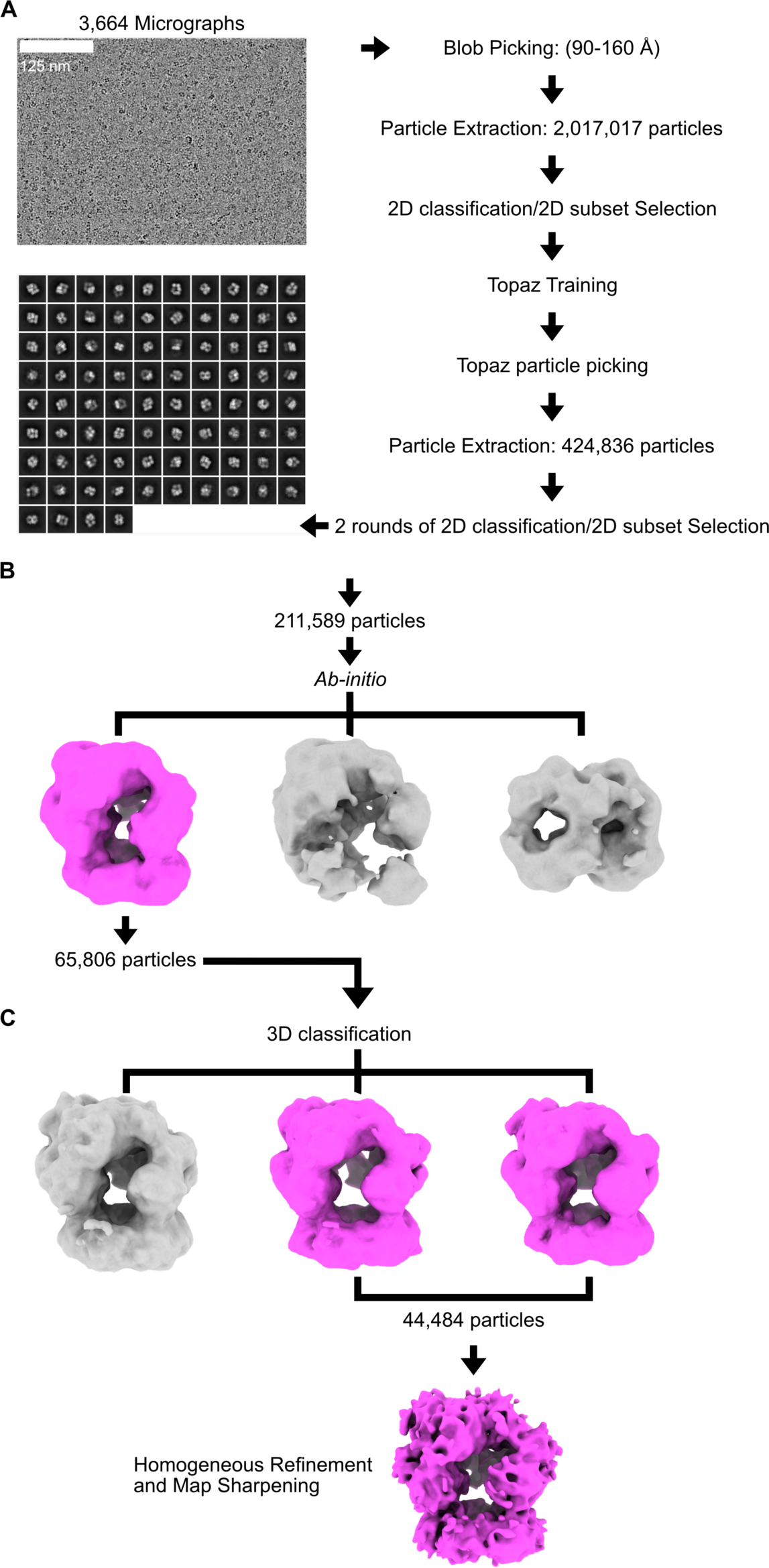
Schematic of the Cryo-EM processing workflow for the Clamp Loader, Sliding Clamp, ATPγS dataset. All data processing was performed using cryoSPARC. **A)** *Representative aligned micrograph and particle-picking pipeline*. (Left) Particles were first picked with cryoSPARC’s blob-picker tool. Particles were the extracted and 2D classified. Particles from the selected 2D classes were used as templates to train Topaz. Particles picked by Topaz were then extracted and used for downstream processing. (Right) Representative 2D classes following Topaz particle picking. **B)** *Ab-initio reconstruction*. Particles from the selected 2D classes were used to generate three *Ab-initio* models. One of the three models is of the clamp loader/sliding clamp complex (pink) **C)** *3D classfication and reconstruction*. Particles from the selected *ab-initio* class were 3D classfied into three classes. Two of the 3D classes were combined and homogeneous refinement was used to generate the final 3D reconstruction (pink), which was used to build the Initial-Binding model.

**Supplemental Figure 3.**
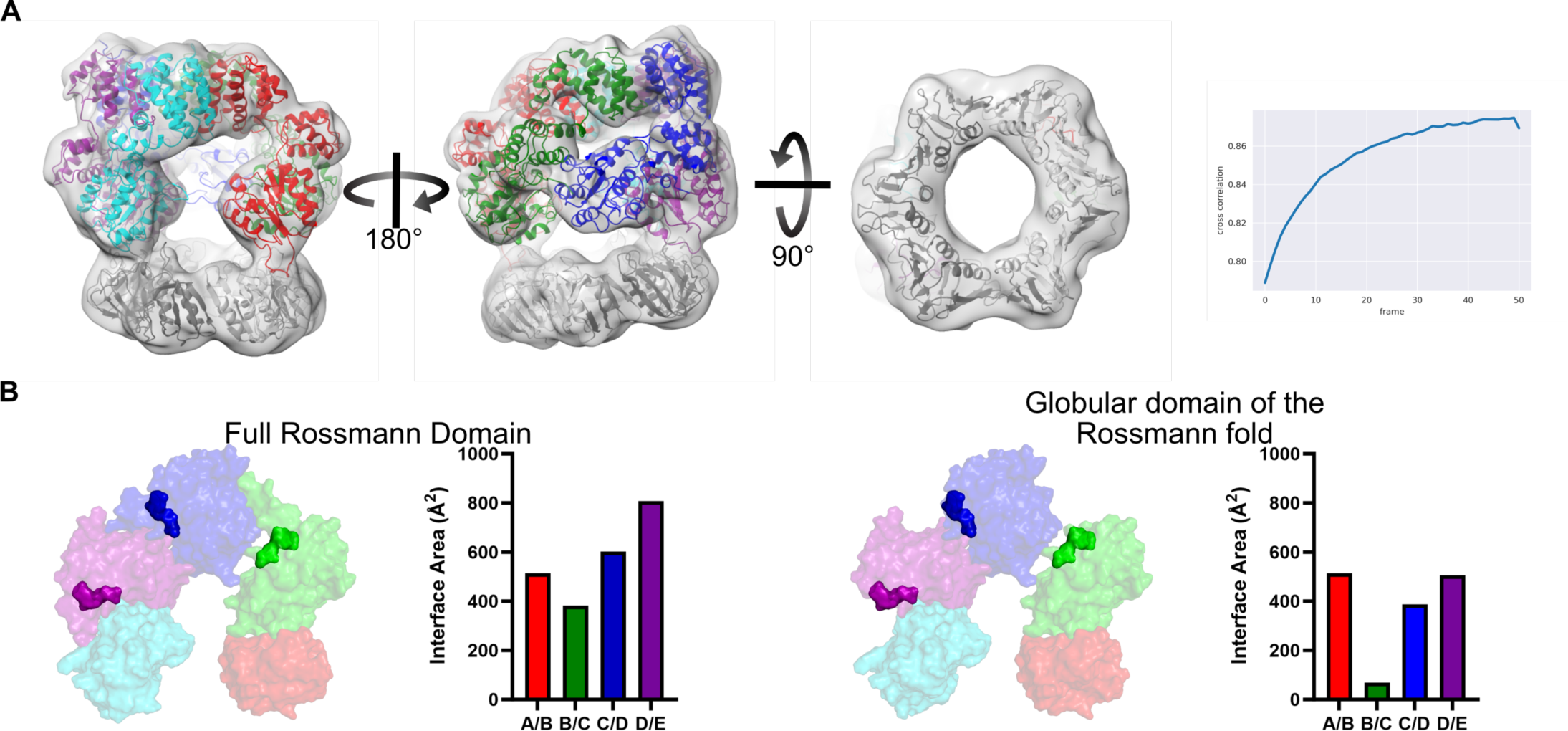
Initial-Binding complex supplemental information. **A)** *Molecular Dynamics Flexible Fitting Model of the Initial-Binding complex*. The Initial-Binding complex model is shown fit into the cryo-EM 3D reconstruction. The Model-to-Map cross-correlation throughout the simulation. Energy minimization was applied during the last iteration which results in a decrease in the overall cross-correlation. **B)** *Geometry of the Rossmann Domain of the Initial-Binding complex*. A transparent surface views of the Rossmann domain of the AAA+ module (left) and globular region of the Rossmann domain (truncating residues 1-10; right) are shown. ATPγS molecules are modelled into the ATP binding sites and shown as solid surfaces. The contact area between adjacent subunits are quantified to the right of each model.

**Supplemental Figure 4.**
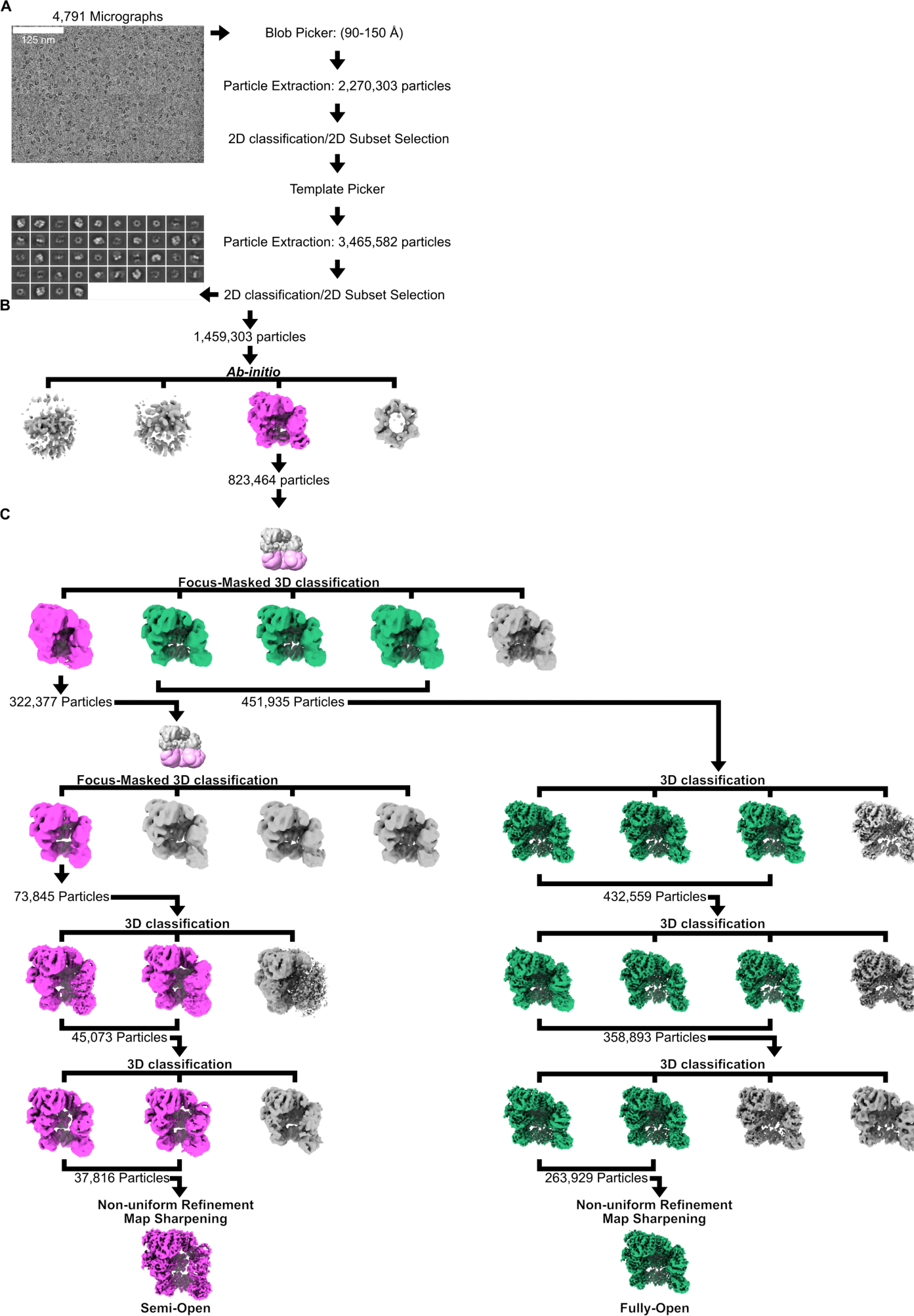
Schematic of the Cryo-EM processing workflow for the Clamp Loader, Sliding Clamp, p/t-DNA, and ADP•BeF_x_ dataset. All data processing was performed using CryoSPARC. **A)** *A representative patch-motion corrected micrograph and particle-picking pipeline*. Particles were first picked with CryoSPARC’s blob-picker tool. Identified particles were extracted and 2D classified. Particles from the selected 2D classes were used as templates for CryoSPARC’s template picker. Particles identified by the template picker were extracted and 2D classified. Representative 2D classes were selected following template picking. **B)** *Ab-initio reconstruction*. Particles from the selected 2D classes were used to generate four *Ab-initio* models. One of the four models resulted in a reconstruction of the clamp loader/sliding clamp (pink). **C)** *3D classification and reconstruction*. A focus mask was applied to the sliding clamp of the *ab-initio* model, which was used to perform a focused 3D classification of the particles into five classes. One of the resulting classes was of a “semi-open” clamp loader/sliding clamp complex (pink), while three were of the open clamp loader/sliding clamp complex (green). Another focused 3D classification was performed on this “Semi-Open” class using a mask on the sliding clamp. One of the resulting classes (pink) was selected for further 3D classification. Non-uniform refinement and map sharpening was used on the final particle stack. This 3D recontruction was used to build the Semi-Open model. The three open classes (green, right) were combined and further 3D classified. Non-uniform refinement and map sharpening was used on the final particle stack. This 3D recontruction was used to build the Fully-Open model.

**Supplemental Figure 5.**
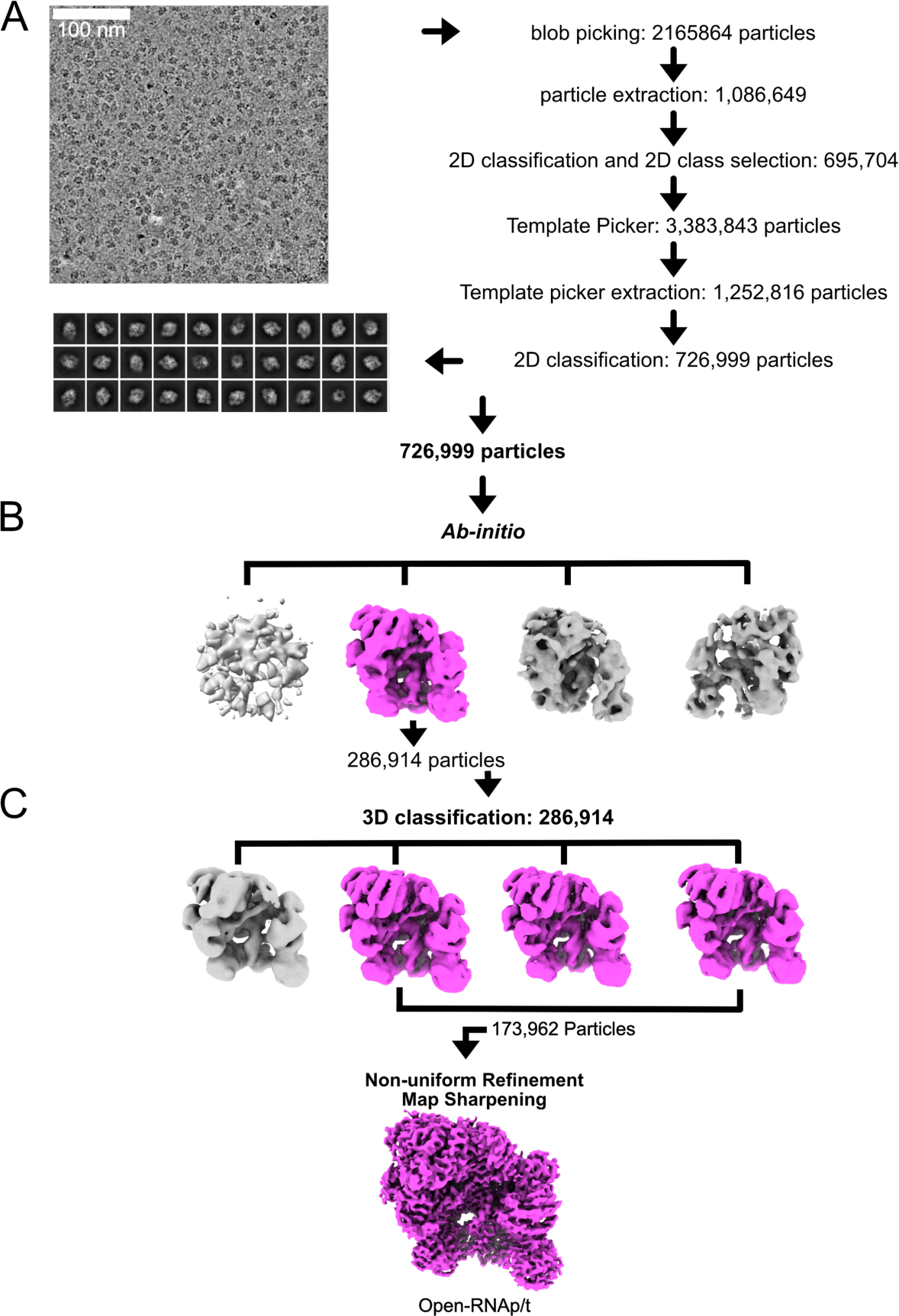
Schematic of the Cryo-EM processing workflow for the Clamp Loader, Sliding Clamp, p/t-RNA, and ADP•BeF_x_ dataset. All data processing was performed using CryoSPARC. **A)** *A representative patch-motion corrected micrograph and particle-picking pipeline*. Particles were first picked with CryoSPARC’s blob-picker tool. Identified particles were extracted and 2D classified. Particles from the selected 2D classes were used as templates for CryoSPARC’s template picker. Particles identified by the template picker were extracted and 2D classified. Representative 2D classes were selected following template picking. **B)** *Ab-initio reconstruction*. Particles from the selected 2D classes were used to generate four *Ab-initio* models. One of the four models resulted in a reconstruction of the clamp loader/sliding clamp (pink). **C)** *3D classification and reconstruction*. The selected particles were 3D classified into four classes. Non-uniform refinement and map sharpening was used on the final particle stack. This 3D recontruction was used to build the Open-RNAp/t model.

**Supplemental Figure 6.**
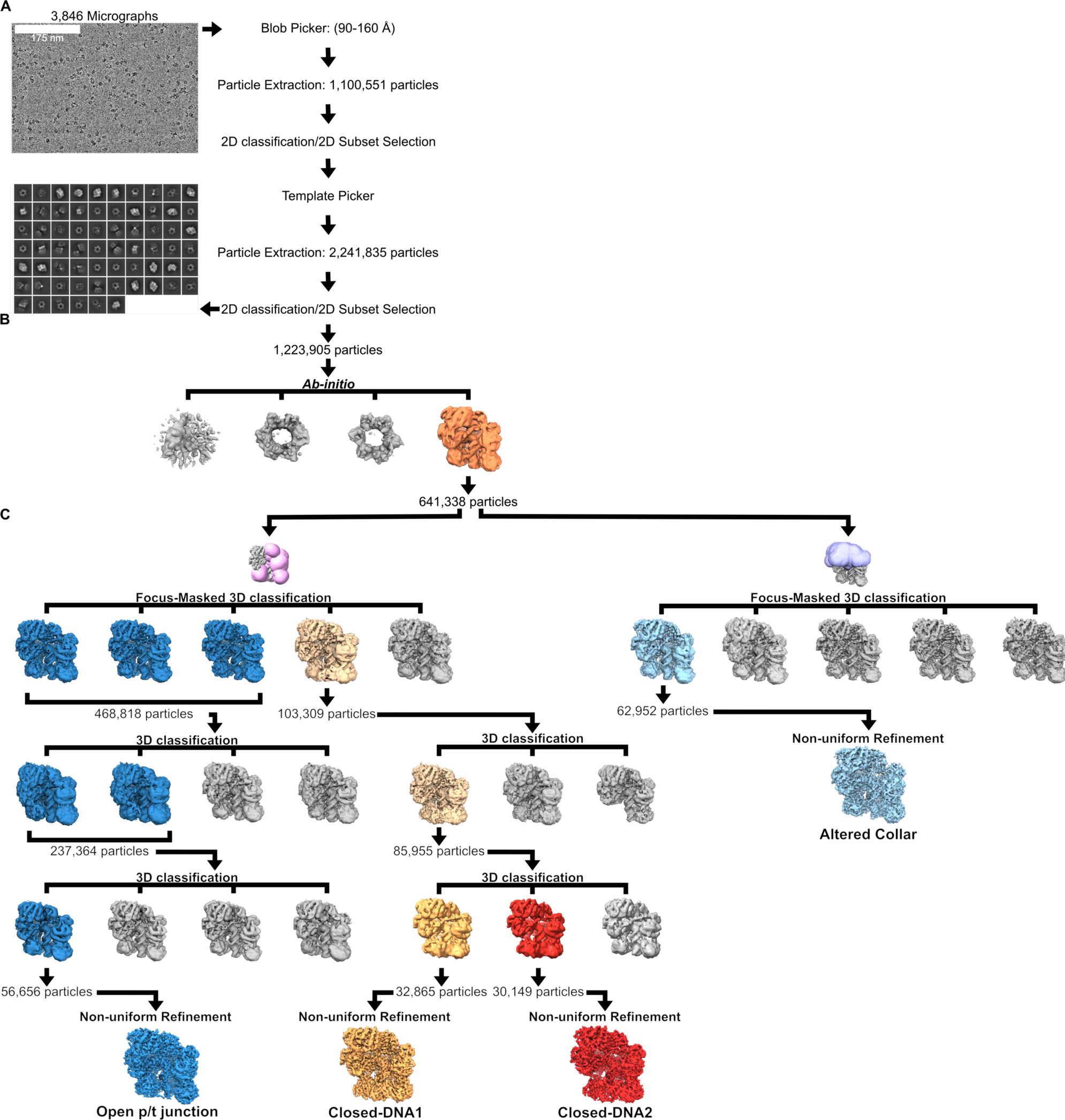
Schematic of the Cryo-EM processing workflow for the Clamp Loader, Sliding Clamp, p/t-DNA, and ADP•BeF_x_ dataset. All data processing was performed using CryoSPARC. **A)** *A representative patch-motion corrected micrograph and particle-picking pipeline*. Particles were first picked with CryoSPARC’s blob-picker tool, extracted, and 2D classified. Particles from the selected 2D classes were used as templates for CryoSPARC’s template picker. Particles identified by the template picker were extracted and 2D classified. **B)** *Ab-initio reconstruction*. Particles from the selected 2D classes were used to generate four *Ab-initio* models. One of the four models consisted of the clamp loader/sliding clamp/p/t-DNA complex (orange) **C)** *3D classification and reconstruction*. The first round of 3D classification was performed by applying a focus mask to the sliding clamp and the A and B subunits of the clamp loader to generate five classes. This resulted in three open clamp loader/sliding clamp/p/t-DNA complexes (blue) and one closed clamp loader/sliding clamp complex (tan). The three clamp loader/sliding clamp/p/t-DNA complexes were pooled and iteratively 3D classified. The final class had density for the clamp loader, p/t-DNA, and all six domains of the sliding clamp. The closed class was 3D classified resulting in two classes of the closed clamp loader/sliding clamp/DNAp/t complex (orange and red). After 3D classification, non-uniform refinement was performed to generate the final reconstructions of the Open-p/t-DNA, Closed-DNA1, and Closed-DNA2 complexes. In parallel, the particles from the *ab-initio* model were 3D classified into five classes while appling a focus mask to the collar and lid domains of the clamp loader/sliding clamp/DNAp/t complex (right). One of the classes had an altered collar (cyan). After 3D classification, non-uniform refinement was performed to generate the final reconstruction of the Altered-Collar class.

**Supplemental Figure 7.**
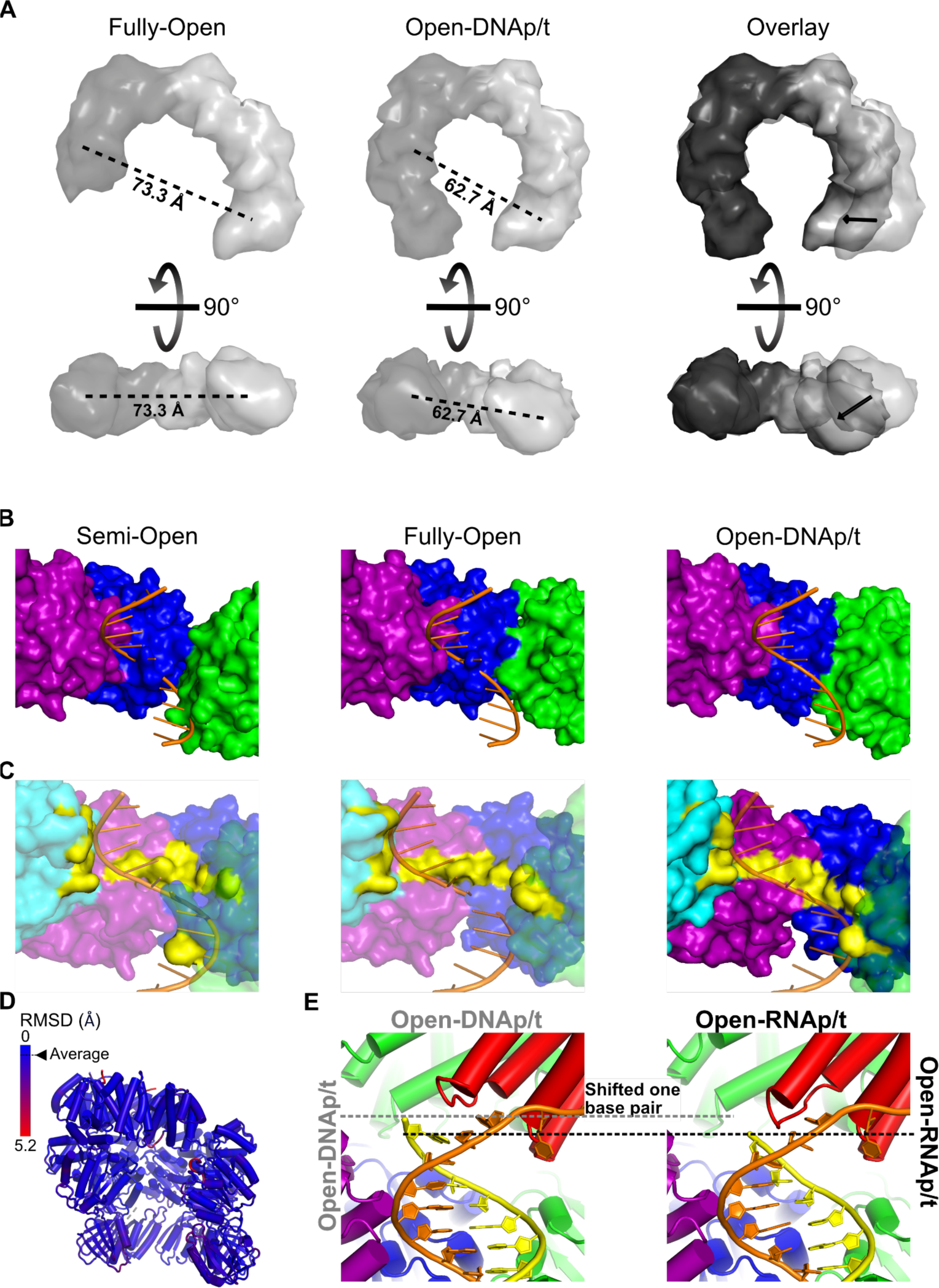
Conformational change induced by p/t-junction binding. **A)** *Conformational changes of the sliding clamp between states*. Models of the sliding clamp of the Fully-Open and Open-DNAp/t states are displayed as low contour surfaces. The distance between domain III of subunit I and domain II of subunit of II of the sliding clamp in each state is shown. The Fully-Open and Open-DNAp/t sliding clamps were aligned on subunit II and displacements of domains within subunit I are shown as vectors. **B)** *Compatibility of the Rossmann domain with template DNA binding*. Surfaces of the Rossmann domains of the B-D subunits of the Semi-Open, Fully-Open, and Open-DNAp/t models are shown. Models were aligned on the D subunit and the template strand from the Open-DNAp/t model is superimposed on the Semi-Open and Fully-Open models. **C)** *Compatibility of template binding residues with template DNA binding*. Surfaces of the Rossmann domains of the B-E subunits of the Semi-Open, Fully-Open, and Open-DNAp/t models are shown. Models were aligned on the E subunit and the template strand from the Open-DNAp/t model is superimposed on the Semi-Open and Fully-Open models. Key template strand gripping residues are highlighted in yellow. **D)** Per residue RMSD of the clamp loader/sliding clamp complex comparing the Open DNAp/t and Open-RNAp/t states. Structures were globally aligned and per-residue C_α_ RMSD was calculated. **E)** *Alignment of template DNA binding residues with the template strand in different states*. Surfaces are shown of the Rossmann domains of the B-E subunits of the Semi-Open, Fully-Open, and Open-DNAp/t models. Models were aligned on the E subunit and the template strand from the Open-DNAp/t model is superimposed on the Semi-Open and Fully-Open models. Residues that bind to the template strand are shown in yellow.

**Supplemental Figure 8.**
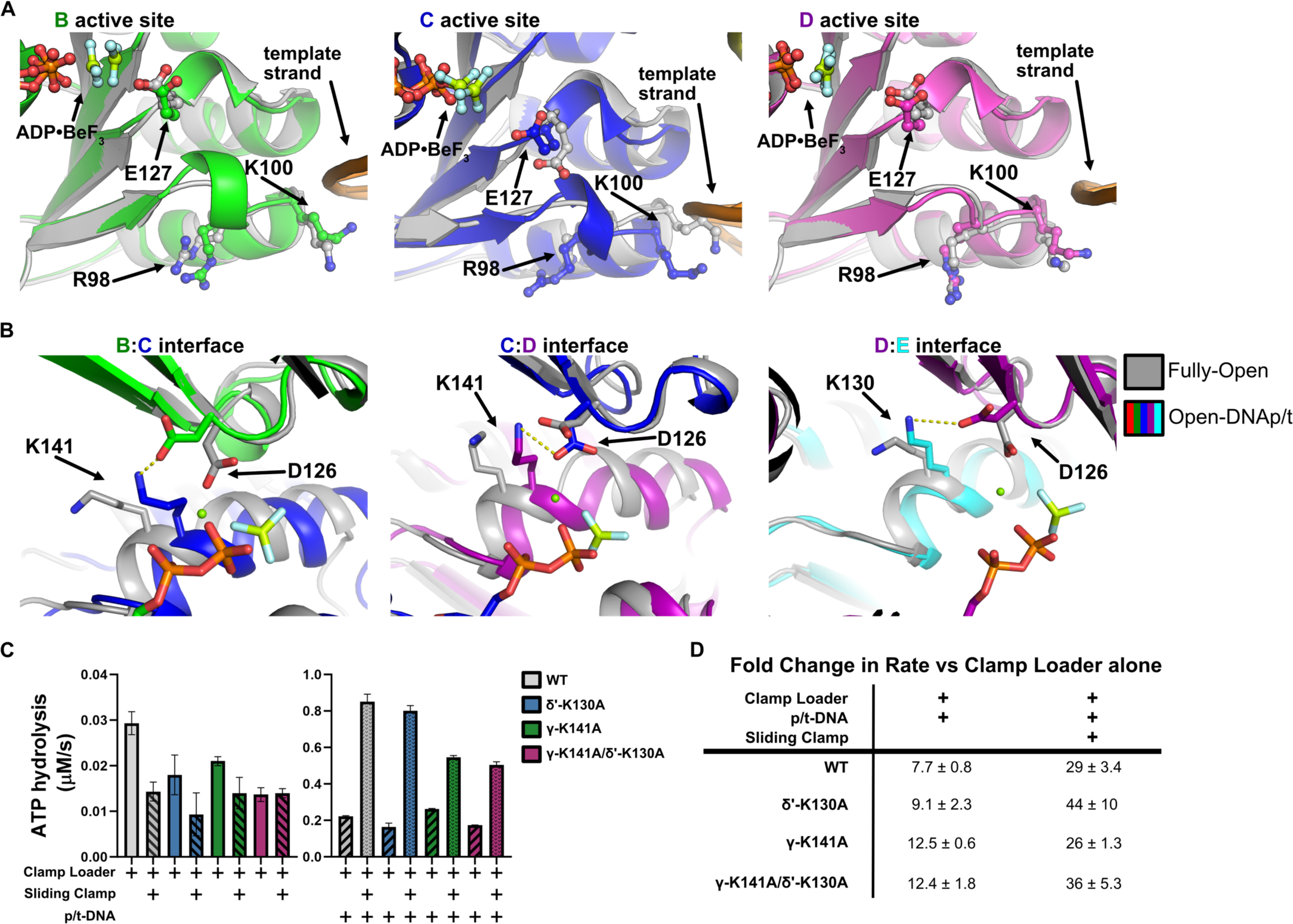
Examination of ATPase activation mechanism. **A)** *No repositioning of the hypothesized “switch” residues*. Overlay of the Fully-Open (gray) and Open-DNAp/t (color) models. In each panel, the models are aligned on the Rossmann domain of the ATP binding subunit. It was hypothesized that R98 and K100 act as allosteric switches by controlling the conformation of the catalytic residue E127 in response to p/t binding^3,16^. No such repositioning is observed. **B)** *Repositioning of the hypothesized DNA-activating lysines*. Overlay of the Fully-Open (gray) and Open-DNAp/t (color) models. In each panel, the models are aligned on the Rossmann domain of the ATP binding subunit of the interface. The *trans*-acting lysine and *cis*-aspartate 126 are shown as sticks. **C)** *The steady-state ATP hydrolysis rate of the Wild-Type*, δ’-*K130A, γ-K141A, and γ-K141A/*δ’-*K130A clamp loaders*. The steady-state ATP hydrolysis rates of the clamp loaders alone and in response to β-clamp (left). The steady-state ATP hydrolysis rate of the clamp loaders in response to p/t-DNA or p/t-DNA and β-clamp (right). **D**) *Fold change of the ATP hydrolysis rate in response to p/t-DNA*. The fold change ± standard deviation of the ATP hydrolysis rate of the clamp loader + DNAp/t or clamp loader + sliding clamp and DNAp/t over the clamp loader alone is shown.

**Supplemental Table 1.**
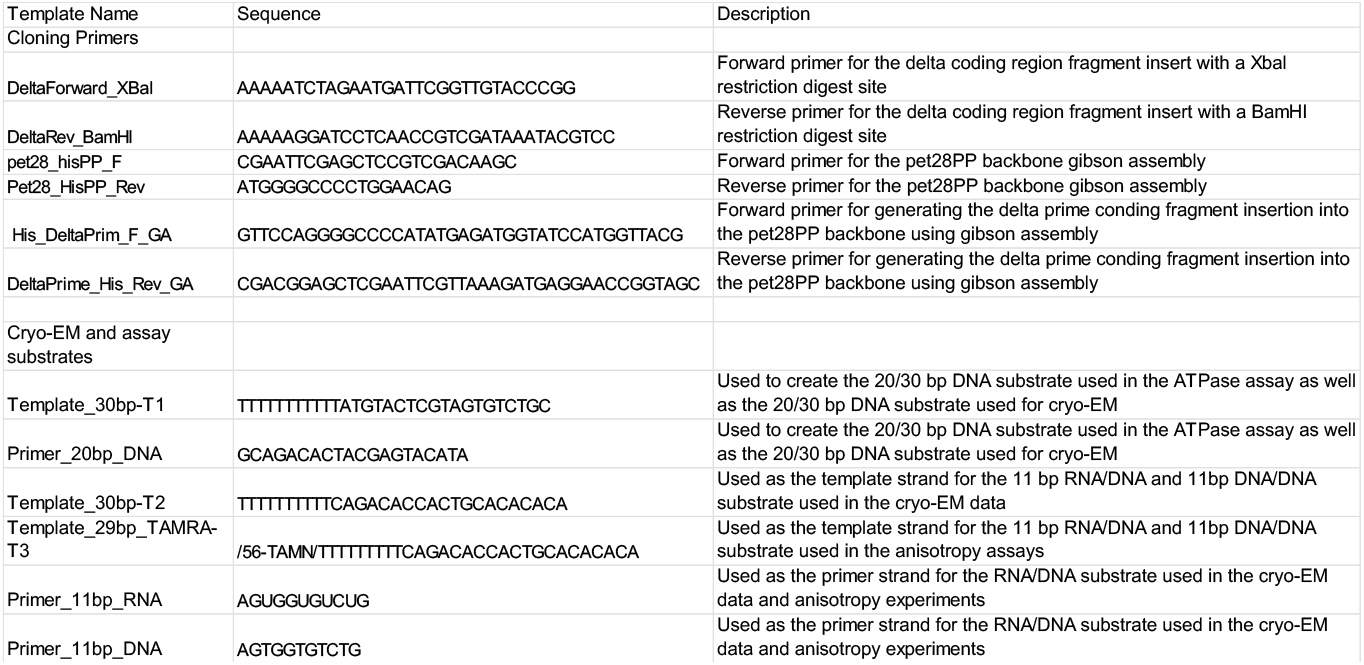
Oligonucleotides used in this study.

**Supplemental Table 2.**
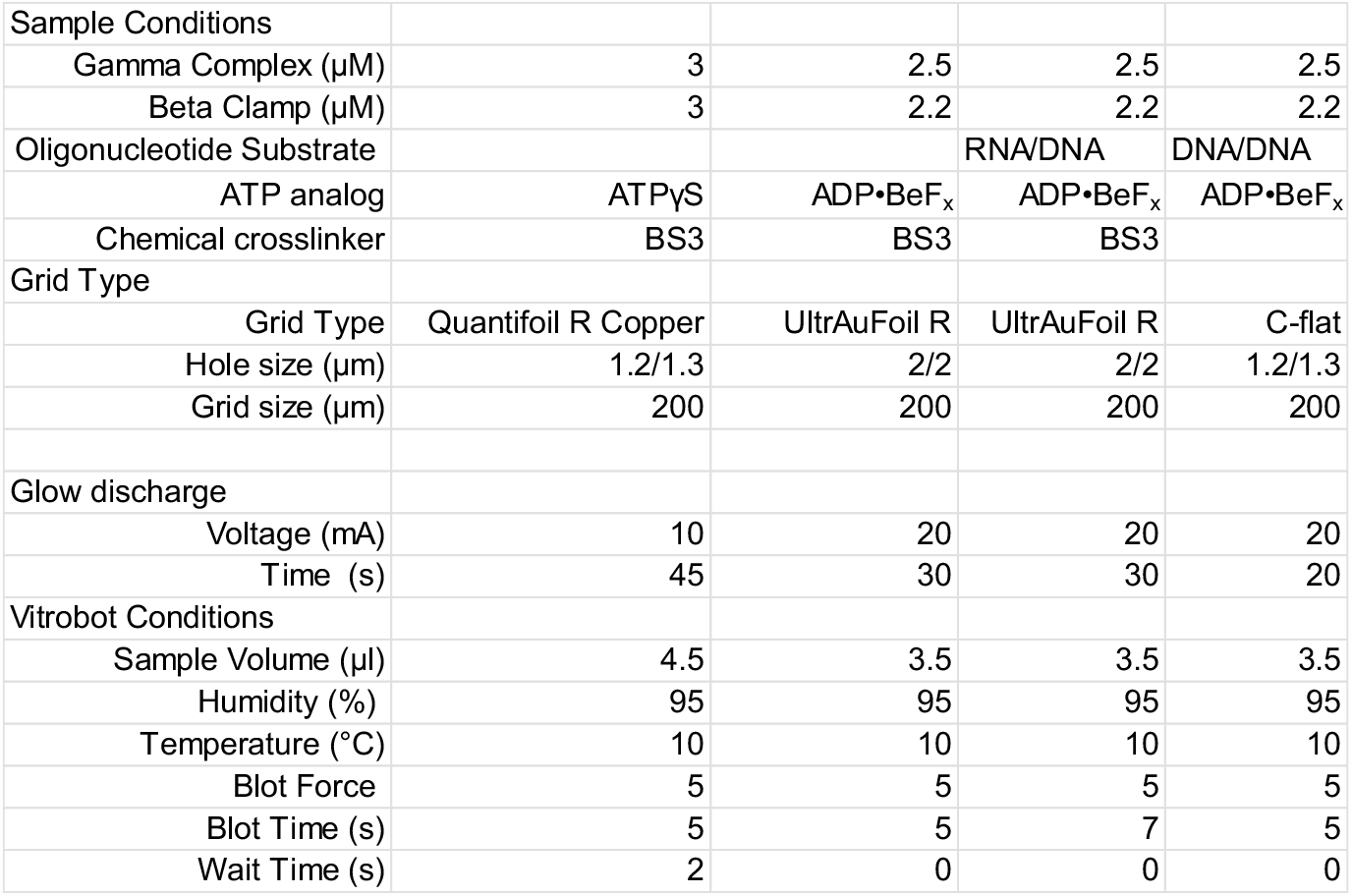
Cryo-EM sample preparation conditions.

|  |  |  |  |  |
| --- | --- | --- | --- | --- |
| Sample Conditions |  |  |  |  |
| Gamma Complex (μM) | 3 | 2.5 | 2.5 | 2.5 |
| Beta Clamp (μM) | 3 | 2.2 | 2.2 | 2.2 |
| Oligonucleotide Substrate |  |  | RNA/DNA | DNA/DNA |
| ATP analog | ATPγS | ADP•BeF<sub>x</sub> | ADP•BeF<sub>x</sub> | ADP•BeF<sub>x</sub> |
| Chemical crosslinker | BS3 | BS3 | BS3 |  |
| Grid Type |  |  |  |  |
| Grid Type | Quantifoil R Copper | UltrAuFoil R | UltrAuFoil R | C-flat |
| Hole size (μm) | 1.2/1.3 | 2/2 | 2/2 | 1.2/1.3 |
| Grid size (μm) | 200 | 200 | 200 | 200 |
| Glow discharge |  |  |  |  |
| Voltage (mA) | 10 | 20 | 20 | 20 |
| Time (s) | 45 | 30 | 30 | 20 |
| Vitrobot Conditions |  |  |  |  |
| Sample Volume (μl) | 4.5 | 3.5 | 3.5 | 3.5 |
| Humidity (%) | 95 | 95 | 95 | 95 |
| Temperature (°C) | 10 | 10 | 10 | 10 |
| Blot Force | 5 | 5 | 5 | 5 |
| Blot Time (s) | 5 | 5 | 7 | 5 |
| Wait Time (s) | 2 | 0 | 0 | 0 |

**Supplemental Table 3.**
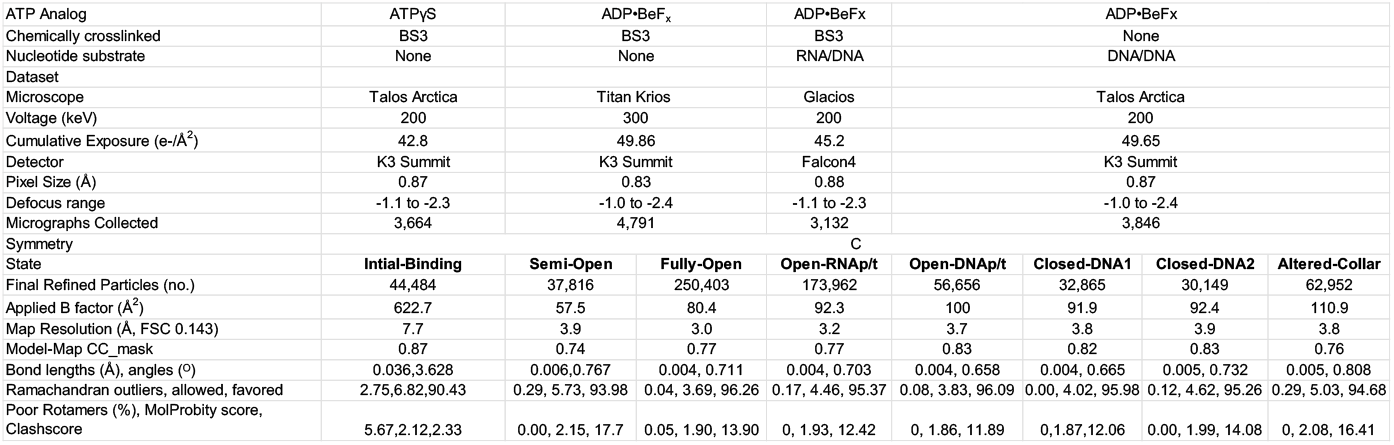
Cryo-EM data collection, processing, and model statistics.

